# mRNA-1273 or mRNA-Omicron boost in vaccinated macaques elicits comparable B cell expansion, neutralizing antibodies and protection against Omicron

**DOI:** 10.1101/2022.02.03.479037

**Authors:** Matthew Gagne, Juan I. Moliva, Kathryn E. Foulds, Shayne F. Andrew, Barbara J. Flynn, Anne P. Werner, Danielle A. Wagner, I-Ting Teng, Bob C. Lin, Christopher Moore, Nazaire Jean-Baptiste, Robin Carroll, Stephanie L. Foster, Mit Patel, Madison Ellis, Venkata-Viswanadh Edara, Nahara Vargas Maldonado, Mahnaz Minai, Lauren McCormick, Christopher Cole Honeycutt, Bianca M. Nagata, Kevin W. Bock, Caitlyn N. M. Dulan, Jamilet Cordon, John-Paul M. Todd, Elizabeth McCarthy, Laurent Pessaint, Alex Van Ry, Brandon Narvaez, Daniel Valentin, Anthony Cook, Alan Dodson, Katelyn Steingrebe, Dillon R. Flebbe, Saule T. Nurmukhambetova, Sucheta Godbole, Amy R. Henry, Farida Laboune, Jesmine Roberts-Torres, Cynthia G. Lorang, Shivani Amin, Jessica Trost, Mursal Naisan, Manjula Basappa, Jacquelyn Willis, Lingshu Wang, Wei Shi, Nicole A. Doria-Rose, Adam S. Olia, Cuiping Liu, Darcy R. Harris, Andrea Carfi, John R. Mascola, Peter D. Kwong, Darin K. Edwards, Hanne Andersen, Mark G. Lewis, Kizzmekia S. Corbett, Martha C. Nason, Adrian B. McDermott, Mehul S. Suthar, Ian N. Moore, Mario Roederer, Nancy J. Sullivan, Daniel C. Douek, Robert A. Seder

## Abstract

SARS-CoV-2 Omicron is highly transmissible and has substantial resistance to antibody neutralization following immunization with ancestral spike-matched vaccines. It is unclear whether boosting with Omicron-specific vaccines would enhance immunity and protection. Here, nonhuman primates that received mRNA-1273 at weeks 0 and 4 were boosted at week 41 with mRNA-1273 or mRNA-Omicron. Neutralizing antibody titers against D614G were 4760 and 270 reciprocal ID_50_ at week 6 (peak) and week 41 (pre-boost), respectively, and 320 and 110 for Omicron. Two weeks after boost, titers against D614G and Omicron increased to 5360 and 2980, respectively, for mRNA-1273 and 2670 and 1930 for mRNA-Omicron. Following either boost, 70-80% of spike-specific B cells were cross-reactive against both WA1 and Omicron. Significant and equivalent control of virus replication in lower airways was observed following either boost. Therefore, an Omicron boost may not provide greater immunity or protection compared to a boost with the current mRNA-1273 vaccine.

## Introduction

The COVID-19 mRNA vaccines BNT162b2 and mRNA-1273 provide highly effective protection against symptomatic and severe infection with ancestral SARS-CoV-2 (Baden et al., 2021b; Dagan et al., 2021; Pilishvili et al., 2021; Polack et al., 2020). More recently, protective efficacy has declined due to both waning vaccine-elicited immunity (Baden et al., 2021a; Bergwerk et al., 2021; Goldberg et al., 2021) and antigenic shifts in variants of concern (VOC) including B.1.351 (Beta) and B.1.617.2 (Delta) (Planas et al., 2021; Wang et al., 2021a; Wang et al., 2021b). Importantly, the introduction of a boost after the initial vaccine regimen enhances immunity and vaccine efficacy against symptomatic disease, hospitalization and death across a broad range of age groups (Andrews et al., 2022; Bar-On et al., 2021; Barda et al., 2021; Garcia-Beltran et al., 2022; Pajon et al., 2022). However, the timing and selection of a boost is a major scientific and clinical challenge during this evolving pandemic in which emerging VOC have distinctive patterns of transmission and virulence and against which vaccine-elicited antibody neutralization is reduced.

The most recent VOC, B.1.1.529, henceforth referred to by its WHO designation of Omicron, was first identified in South Africa in November 2021 and was associated with a dramatic increase in COVID-19 cases (Cele et al., 2021; Maslo et al., 2021). Omicron is highly contagious, with a significant transmission advantage compared to Delta, which until recently was the dominant VOC worldwide (Viana et al., 2022). It remains unclear, however, if this advantage is due to differences in cell entry, enrichment in respiratory aerosols, or the ability to evade immunity conferred by vaccination or prior infection. Compared to the ancestral strain, Omicron contains more than 30 mutations in the spike (S) gene, including S477N, T478K, E484A, Q493R, G496S, Q498R, N501Y and Y505H in the receptor binding motif (RBM) alone. Neutralizing antibody titers in sera of individuals recently recovered from previous infection or shortly after immunization with two doses of an mRNA-based COVID-19 vaccine are dramatically reduced to Omicron as compared to the ancestral strains Wuhan-Hu-1, USA-WA1/2020 (WA1) and D614G. Numerous studies using both live virus and pseudovirus neutralization assays report a 60- to 80-fold reduction for convalescent sera and a 20- to 130-fold reduction for vaccinee sera (Edara et al., 2021a; Hoffmann et al., 2021; Muik et al., 2022; Schmidt et al., 2021). mRNA-1273 vaccine efficacy against breakthrough cases of Omicron in the first few months after immunization has been estimated as 30% in California, USA and 37% in Denmark (Hansen et al., 2021; Tseng et al., 2022) and a complete loss of protection within six months (Accorsi et al., 2022). Multiple reports have suggested that Omicron has reduced virulence compared to prior VOC in humans, mice and hamsters (Davies et al., 2022; Halfmann et al., 2022; Suryawanshi et al., 2022). It is possible that reduced virulence of Omicron may result from preferential replication in the upper airway compared to the lungs, perhaps due to altered cellular tropism not reliant on expression of transmembrane serine protease 2 (TMPRSS2) (Meng et al., 2022; Willett et al., 2022). However, the effect of any reduction in intrinsic viral pathogenicity may be somewhat offset in the context of reduced vaccine efficacy and enhanced virus transmission in human populations worldwide. Together these data reinforce the value of boosting to limit the extent of infection from Omicron.

Variant-matched boosts have been suggested as a strategy to enhance neutralizing and binding antibody titers to the corresponding VOC beyond the levels conferred by existing FDA-approved boosts, which are homologous to the original ancestral WA1-matched primary vaccine regimen. We previously showed that boosting mRNA-1273 immunized nonhuman primates (NHP) with mRNA-1273 or a boost matched to the Beta VOC spike (mRNA-1273.351 or mRNA-1273.Beta) resulted in significant enhancement of neutralizing antibody responses across all VOC tested and an expansion of S-specific memory B cells with ∼80-90% expressing WA1 and Beta. Moreover, both vaccines provided substantial and similar protection against Beta replication (Corbett et al., 2021a). These NHP data were confirmed in a study in humans which compared an mRNA-1273 boost to mRNA-1273.Beta ∼6 months after participants had received the standard two-dose mRNA-1273 vaccine regimen (Choi et al., 2021). Following either boost, neutralizing titers were substantially increased against D614G and several variants including Beta and were comparable between boost groups. Of note, the level of neutralizing antibodies to Beta after either boost were about 10-fold higher than after the initial vaccination suggesting affinity maturation or epitope focusing of the B cell response. Together, these data suggest that the variant Beta boost did not uniquely enhance immunity or protection compared to existing ancestral strain-matched boosts. However, as Omicron contains more mutations in S compared to Beta and demonstrates even more substantial escape from vaccine-elicited neutralizing antibodies than does Beta, it is unclear whether an Omicron-specific boost would provide an additional protective benefit against Omicron infection beyond that of WA1-matched boosts.

The nonhuman primate (NHP) model has been useful for demonstrating immune correlates, mechanisms and durability of vaccine-elicited protection against SARS-CoV-2 and been largely predictive for what has been observed in humans in terms of protective efficacy (Corbett et al., 2021b; Gagne et al., 2022; Gilbert et al., 2021). Here, we vaccinated NHP with 100µg mRNA-1273 at weeks 0 and 4, which is a similar dose and schedule as used in humans. Animals were then boosted about ∼9 months later with 50µg of either a homologous dose of mRNA-1273 or mRNA-1273.529, which is matched to Omicron S (henceforth referred to as mRNA-Omicron). For the duration of these 9 months, we collected sera, bronchoalveolar lavage (BAL) and nasal washes to analyze the kinetics of antibody binding and neutralization as well as the frequency of S-specific B cells for WA1 and Omicron as well as Beta and Delta. Four weeks after boost, NHP were challenged with Omicron. Viral replication in upper and lower airways and lung inflammation were measured to compare boost-elicited protection against Omicron.

## Results

### Kinetics of serum antibody responses following mRNA-1273 immunization and boost

Indian-origin rhesus macaques (*n*=8) were immunized with 100µg of mRNA-1273 at weeks 0 and 4 (Fig. S1). Sera were collected at weeks 6 (peak) and 41 (memory) to measure immunoglobulin G (IgG) binding to WA1 S and a panel of VOC (Fig. 1A). At week 6, we observed a clear hierarchy of binding titers with WA1>Delta>Beta >Omicron. Geometric mean titers (GMT) to WA1 and Omicron were 8x10^19^ and 3x10^15^ area under the curve (AUC). Antibody titers waned markedly by week 41, with GMT of 2x10^12^ and 2x10^8^ AUC for WA1 and Omicron, reflecting a 7-log decline for each strain. Similar antibody kinetics and hierarchy of potency were observed when measuring binding to the receptor binding domain (RBD) of the same variants, with titers to Omicron of 7x10^11^ AUC at week 6 and 8x10^7^ AUC at week 41 (Fig. 1B). Nine months after the second dose of mRNA-1273 (week 41), NHP were boosted with 50µg of homologous mRNA-1273 or heterologous virus challenge-matched mRNA-Omicron (*n*=4/group). S-binding titers were restored to the same level as observed at week 6 following either a homologous or heterologous boost, and titers to Omicron were still lower than all other variants.

**Figure 1.**
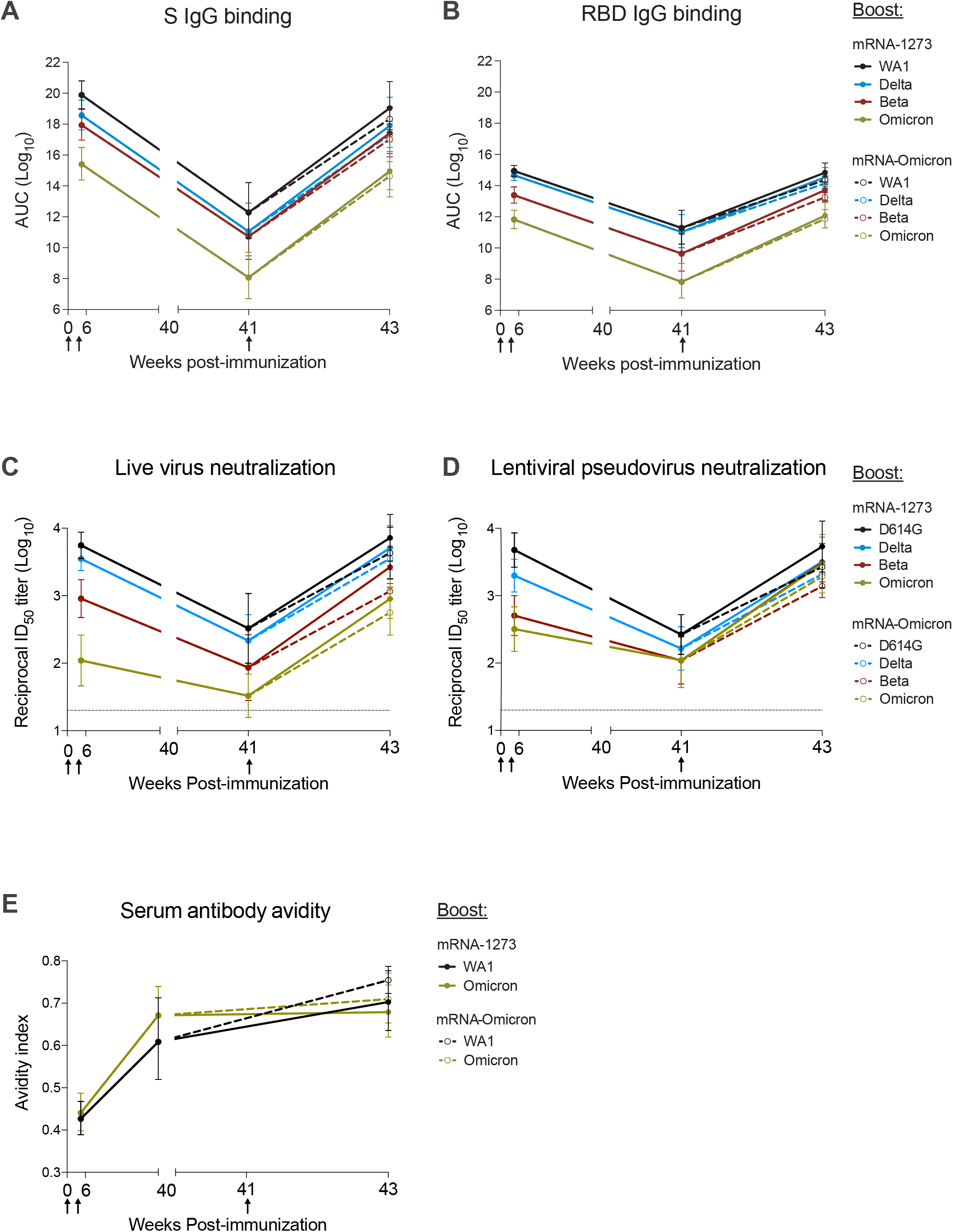
Kinetics of serum antibody responses following mRNA-1273 immunization and boost. (A-E) Sera were collected at weeks 6, 40 or 41, and 43 post-immunization. (A-B) IgG binding titers to (A) variant S and (B) variant RBD expressed in AUC. (C-D) Neutralizing titers to (C) live virus and (D) lentiviral pseudovirus expressed as the reciprocal ID50. (E) Avidity index for WA1 S-2P- and Omicron S-2P-binding serum antibodies. Circles represent geometric means (A-D) or arithmetic means (E). Error bars represent 95% confidence intervals (CI). Assay LOD indicated by dotted lines. Break in X-axis indicates a change in scale without a break in the range depicted. Responses to variants are color-coded as WA1 or D614G (black), Delta (blue), Beta (red) and Omicron (green). Arrows represent timepoints of immunizations. Following the boost at week 41, mRNA-1273-boosted NHP indicated by solid lines and mRNA-Omicron-boosted NHP indicated by dashed lines. 4 NHP per boost group. See also Figure S1 for experimental schema and Table S1 for detailed neutralizing titers.

Neutralizing antibody titers were then assessed using a live virus assay (Fig. 1C and Table S1). At week 6, neutralizing titers were highest to D614G followed by Delta, then Beta and Omicron. Titers to all variants markedly declined by week 41, including a drop in reciprocal 50% inhibitory dilution (ID_50_) titers for D614G from 5560 at week 6 to 330 at week 41 and for Omicron from 110 at week 6 to 33 at week 41. However, following either boost, neutralizing titers to D614G and Delta were increased similar to week 6 and titers to Beta and Omicron were greater than they had been at week 6 (Beta: *P*=0.05 and 0.035; Omicron: *P*=0.041 and 0.01 for mRNA-1273 and mRNA-Omicron, respectively). We substantiated these findings using a lentiviral pseudovirus neutralization assay similar to the one used to assess immune responses in human clinical trials (Fig. 1D). Following either boost, pseudovirus neutralizing titers were greater to Beta and Omicron than they had been at the week 6 time-point, including an increase in Omicron titers from 320 GMT to 2980 GMT in the mRNA-1273 boost group and 1930 GMT in the mRNA-Omicron boost group (Beta: *P*=0.022 and <0.0001; Omicron: *P*=0.049 and 0.002 for mRNA-1273 and mRNA-Omicron, respectively).

The increase in neutralizing titers to all VOC tested after the third dose could suggest continued antibody maturation (Gaebler et al., 2021). To extend this analysis, we measured antibody avidity over time following immunization (Fig. 1E). Serum antibody avidity to WA1 S-2P increased from a geometric mean avidity index of 0.43 to 0.61 from weeks 6 to 41 (*P*=0.0016), a comparable increase to our previous findings (Corbett et al., 2021a; Gagne et al., 2022). Similarly, avidity to Omicron S-2P rose from 0.44 to 0.67 (*P*<0.0001). Following the boost, no further change was observed (*P*>0.05 for both boost groups).

Collectively, these data show that boosting with the homologous mRNA-1273 or mRNA-Omicron leads to comparable and significant increases in neutralizing antibody responses against all VOC including Omicron.

### mRNA-1273 and mRNA-Omicron boosting increase mucosal antibody responses to Omicron

Upper and lower airway antibody response are critical for mediating protection against SARS-CoV-2 and were assessed following immunization. Nasal washes (NW) and bronchoalveolar lavage fluid (BAL) were collected at weeks 8 (four weeks after the initial mRNA-1273 immunizations), 39 (pre-boost) and 43 (two weeks after the boost). At all timepoints, BAL and NW IgG S-binding titers followed the hierarchy of WA1>Delta>Beta>Omicron (Fig. 2A-B), the same trend detected in our serological assays. In BAL, immediately prior to the boost, GMT were 6.8x10^6^, 4.0x10^6^, 1.3x10^6^ and 2.5x10^4^ AUC for WA1, Delta, Beta and Omicron, respectively. These titers correlated with a 2-fold reduction for Delta compared to WA1, a 5-fold reduction for Beta, and a 275-fold reduction for Omicron. Following either boost, titers were increased by 3-4 logs for all variants. In NW, titers decreased from ∼10^11^ for WA1, Delta and Beta at week 8 to 1.3x10^6^, 3.7x10^5^ and 1.9x10^5^ for WA1, Delta and Beta, respectively. GMT to Omicron similarly declined from 8.8x10^8^ to 8.7x10^3^ AUC and were lower than WA1 and all other variants. Consistent with the findings in the BAL, either boost increased nasal antibody titers ∼6-7 logs, with GMT of ∼10^12^ for WA1, Delta and Beta and ∼10^10^ for Omicron.

**Figure 2.**
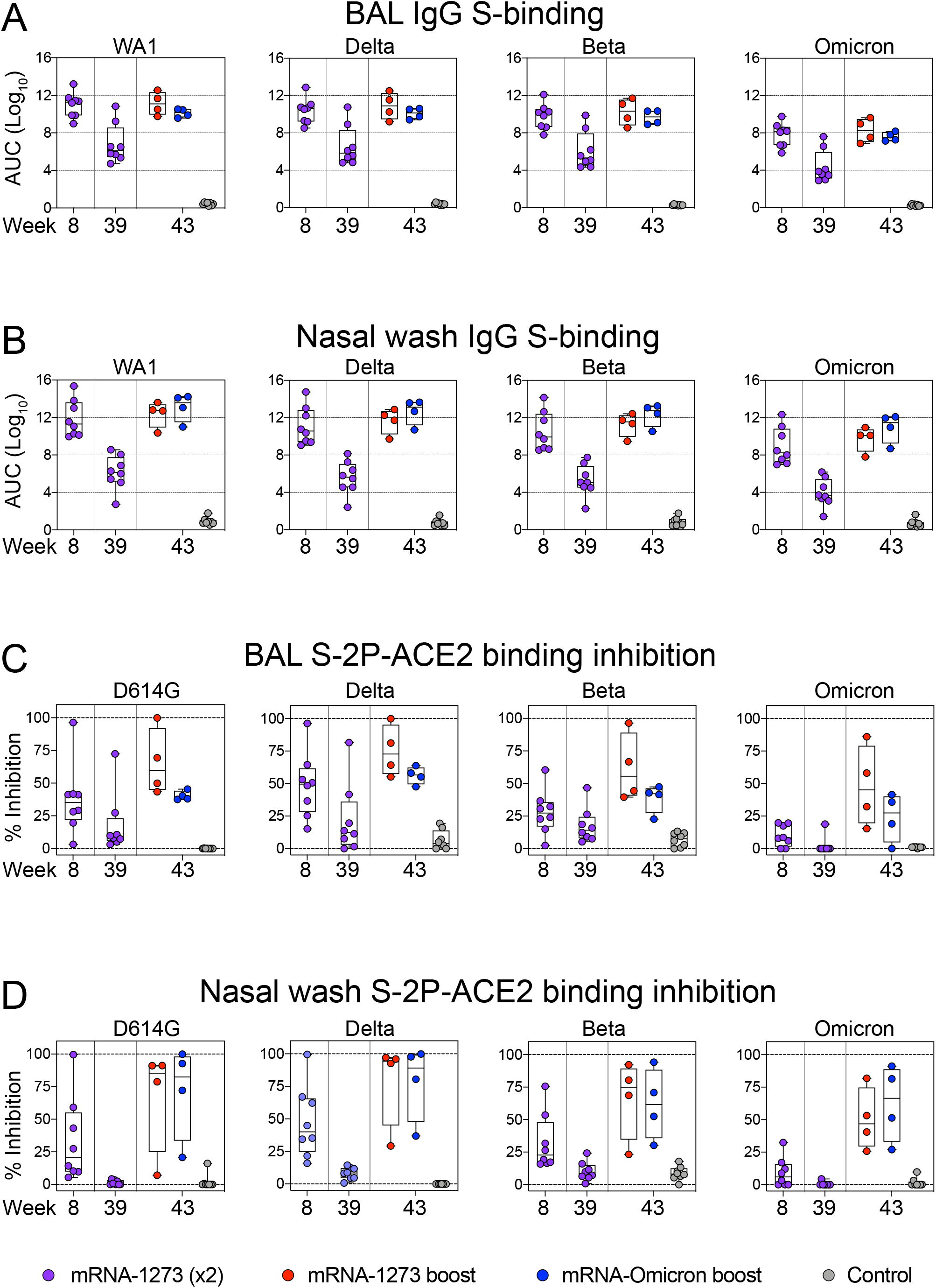
Kinetics of mucosal antibody responses following mRNA-1273 immunization and boost. (A-D) BAL (A, C) and NW (B, D) were collected at weeks 8, 39 and 43 post-immunization. (A-B) IgG binding titers to WA1, Delta, Beta and Omicron expressed in AUC. (C-D) D614G, Delta, Beta and Omicron S-2P-ACE2 binding inhibition in the presence of mucosal fluids. All samples diluted 1:5. Circles indicate individual NHP. Boxes represent interquartile range with the median denoted by a horizontal line. Dotted lines are for visualization purposes and denote 4-log10 increases in binding titers (A-B) or 0 and 100% inhibition (C-D). 8 controls and 8 vaccinated NHP, split into 2 cohorts post-boost. See also Table S2 for list of mutations in variant-specific S-2P-ACE2 inhibition assays.

In a number of prior NHP studies, we have not been able to detect antibody neutralizing titers using pseudo-or live-virus assays from NW or BAL. However, based on its high sensitivity, we have used the angiotensin converting enzyme 2 receptor (ACE2) inhibition assay to measure antibody function as a surrogate for neutralization capacity (Corbett et al., 2021a; Gagne et al., 2022). While the antigen for determination of binding titers was wildtype S, our ACE2 inhibition assay used stabilized S-2P (Table S2). In the BAL, 25-50% median binding inhibition was observed for all variants at week 8, except for Omicron S-2P in which binding inhibition was low to undetectable (Fig. 2C). ACE2 binding inhibition declined to a median of <15% for all variants by week 39 and then increased after either the homologous or mRNA-Omicron boost to levels above those at week 8. Of note, although ACE2 inhibition of Omicron S-2P increased following the boost, it remained lower than all other variants. In the upper airway, ACE2 inhibition was low to undetectable at week 39 following the initial vaccine regimen for all variants. However, after either boost, there was an increase across all variants including Omicron to values higher than the initial peak at week 8 (Fig. 2D). Thus, boosting with either vaccine was important for enhancing mucosal antibody binding and neutralization responses.

### Similar expansion of cross-reactive S-2P-specific memory B cells following boosting

The observation of rapid and significant increases in binding and neutralizing antibody titers to Omicron in both blood and mucosal sites after homologous or heterologous mRNA boost suggests an anamnestic response involving the mobilization of cross-reactive memory B cells. Thus, we measured B cell binding to pairs of fluorochrome-labeled S-2P probes with different VOC including Omicron at weeks 6, 41 and 43 (2 weeks post-boost) (Fig. S2). Of the total S-2P specific memory B cell responses at week 6, 63% were dual-specific and capable of binding both WA1 and Omicron probes, with 33% binding WA1 alone and only 4% which bound Omicron alone (Fig. 3A, 4A). By week 41, the total S-specific memory B cell compartment had declined ∼90% as a fraction of all class-switched memory B cells (Fig. S3A), although the dual-specific population remained the largest fraction within the S-binding pool (Fig. 4A).

**Figure 3.**
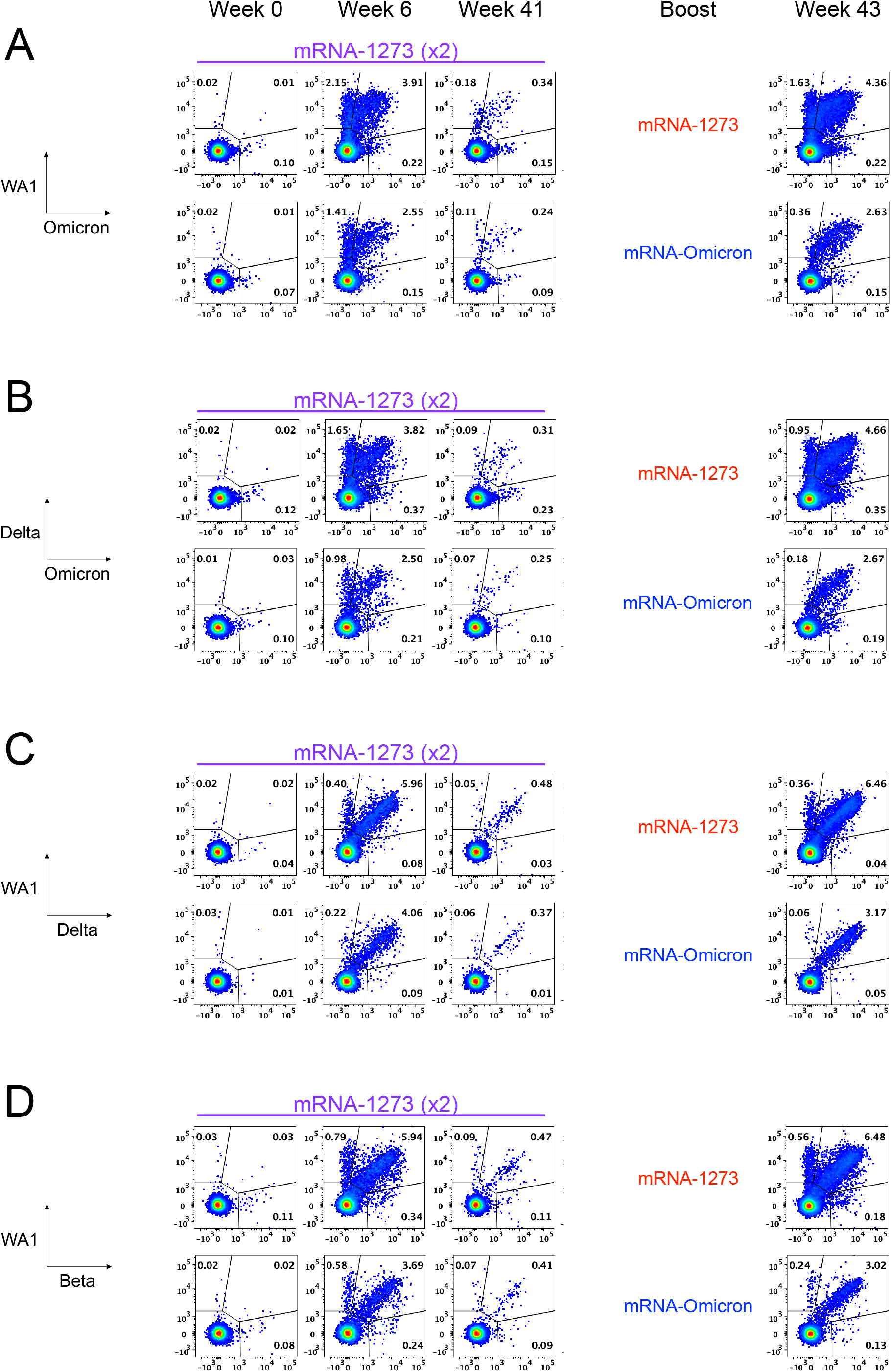
Memory B cell specificities following immunization and boosting. (A-D) Representative flow cytometry plots showing single variant-specific (top left and bottom right quadrant) and dual-variant specific (top right quadrant) memory B cells at weeks 0, 6, 41 and 43 post-immunization. Event frequencies per gate are expressed as a percentage of all class-switched memory B cells. Cross-reactivity shown for (A) WA1 and Omicron S-2P, (B) Delta and Omicron S-2P, (C) WA1 and Delta S-2P and (D) WA1 and Beta S-2P. 4-7 NHP per group. See also Figure S2 for B cell gating strategy.

**Figure 4.**
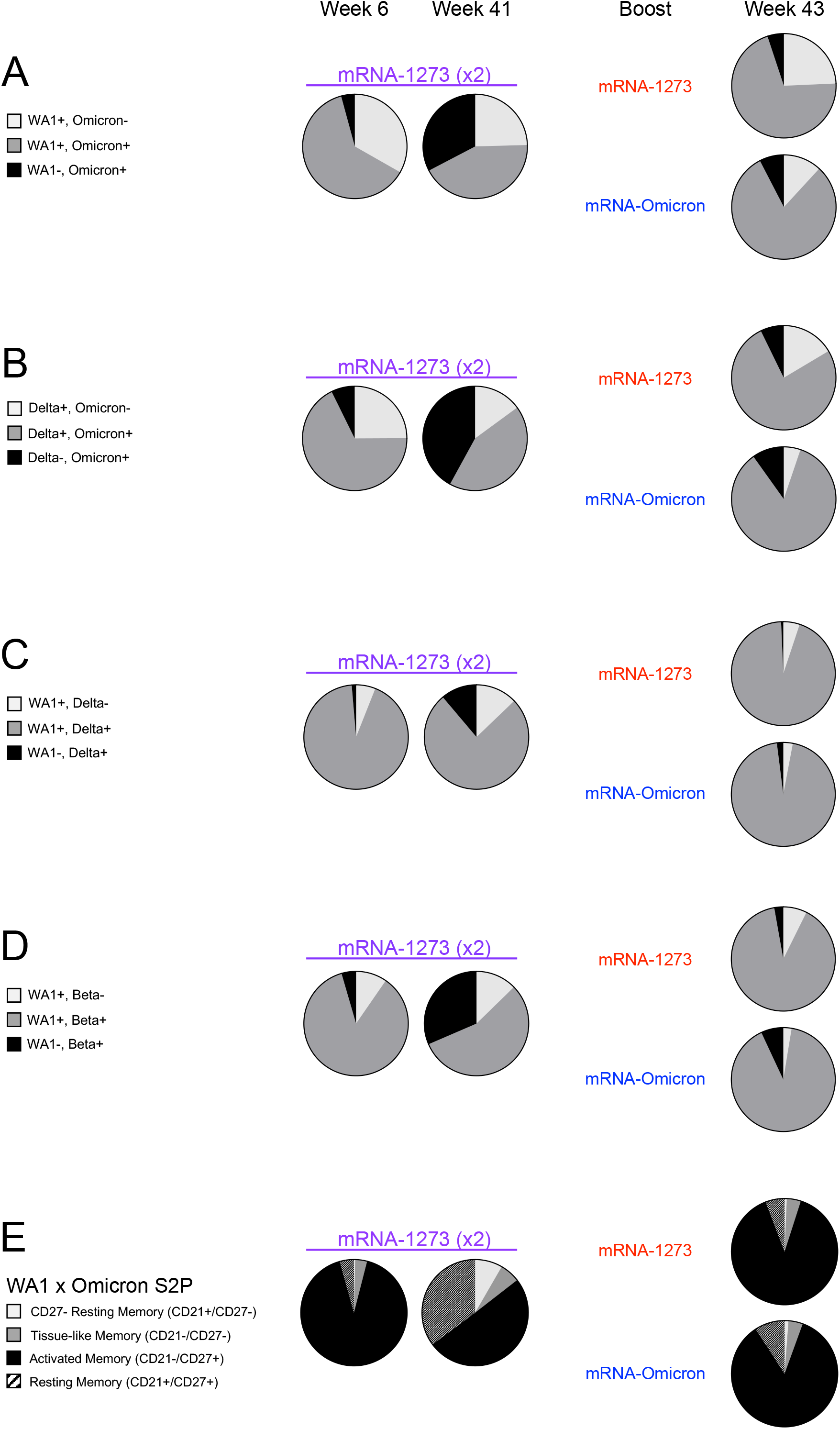
Similar expansion of cross-reactive S-2P-specific memory B cells following boosting. (A-D) Pie charts indicating the proportion of total S-binding memory B cells that are cross-reactive (dark gray) or specific for the indicated variants (black or light gray) for all NHP (geomean) at weeks 6, 41 and 43 post-immunization. Where applicable, memory B cells specific only for WA1 or Delta are represented by the light gray segment. Cross-reactivity shown for (A) WA1 and Omicron S-2P, (B) Delta and Omicron S-2P, (C) WA1 and Delta S-2P and (D) WA1 and Beta S-2P. (E) Pie charts indicating the proportion of total S-2P-binding memory B cells (geomean) that have a phenotype consistent with resting memory (pattern), activated memory (black), tissue-like memory (dark gray) or CD27-negative resting memory (light gray) B cells at weeks 6, 41 and 43 post-immunization. Analysis shown here for memory B cells that bind to WA1 and/or Omicron S-2P. 4-7 NHP per group. See also Figure S3 for frequencies of cross-reactive S-2P memory B cells, Figure S4 for serum epitope reactivity and Figures S4 and S5 for T cell responses after boosting.

Two weeks after boosting, there was an expansion of the total S-specific memory B cell compartment similar to that observed at week 6. Following an mRNA-1273 boost, 24% of all S-2P-specific memory B cells were specific for WA1 alone and 71% were dual-specific for WA1 and Omicron. After the mRNA-Omicron boost, 81% were dual-specific for WA1 and Omicron., with 12% specific for WA1 only (Fig. 4A). Of note, we did not observe a population of Omicron-only memory B cells before or after the boost that was clearly distinct from background staining (Fig. 3A). These data suggest a marked expansion of cross-reactive dual-specific WA1- and Omicron-positive B cells for either boost, with mRNA-1273 also expanding WA1-only B cell responses. The increase in cross-reactive B cells for WA1 and Omicron is consistent with the comparable and high-level of neutralizing titers against D614G and Omicron by either boost (Fig. 1C-D). To extend these data, serologic mapping of antigenic sites on Omicron and WA1 RBD was performed. This analysis revealed that boosting with either mRNA-1273 or mRNA-Omicron elicited serum antibody reactivity with similar RBD specificities (Fig S4).

To further explore the effect of boosting on anamnestic B cell responses, we phenotyped the activation status of S-binding memory B cells (Fig. 4E). WA1 S-2P- and/or Omicron S-2P-binding memory B cells predominantly had an activated memory phenotype immediately after both the second and third doses.

It has recently been shown that individuals in South Africa with or without prior vaccination had increased neutralizing antibody titers to Delta and Omicron following Omicron infection (Khan et al., 2022). Thus, we determined whether there was cross-reactivity of B cells for Delta and Omicron following vaccination. Indeed, 68% of all Delta S-2P and/or Omicron S-2P memory B cells were also dual-specific at week 6 and the remainder of S-binding memory B cells largely bound Delta alone (Fig. 3B, 4B). Following a third dose, the frequency of dual-specific cells increased to 76% for mRNA-1273 and 85% for mRNA-Omicron, consistent with our findings on cross-reactive B cells using WA1 and Omicron S-2P probes.

To complete the analysis and demonstrate cross-reactivity of B cells across other variants, we assessed the frequencies of memory B cells specific for WA-1 and Delta or Beta. We have previously reported that dual-specific WA1 S-2P and Delta S-2P memory B cells accounted for greater than 85% of all memory B cells which bound either spike after two immunizations with mRNA-1273 (Gagne et al., 2022). Here we confirmed and extended these findings and show that after either boost, ∼95% of all WA1- and/or Delta-binding memory B cells are dual-specific (Fig. 3C, 4C). Similar findings were obtained with WA1 and Beta S-2P probes, in which the dual-specific population was 85% at week 6 and 90% following either boost (Fig. 3D, 4D). Of note following the mRNA-Omicron boost, very few B cells were detected that only bound WA1 epitopes when co-staining for Delta or Beta. Overall, the data show that cross-reactive cells were expanded following a boost with either mRNA-1273 or mRNA-Omicron while only mRNA-1273 was capable of boosting memory B cells specific for WA1 alone (Fig. S3).

### S-2P-specific T cell responses in blood and BAL following vaccination

We have previously shown that mRNA-1273 immunization elicits T_H_1, T_FH_ and a low frequency of CD8 responses to S peptides in NHP (Corbett et al., 2020; Corbett et al., 2021a; Corbett et al., 2021c; Gagne et al., 2022; Jackson et al., 2020). Consistent with the prior studies we show that mRNA-1273 elicits T_H_1, T_FH_ and low-level CD8 T cell responses at the peak of the response (week 6) that decline over time (Fig. S5 and S6). Boosting with either mRNA-1273 or mRNA-Omicron increased T_FH_ responses which could be important for expanding the S-specific memory B cell population following the boost (Johnston et al., 2009; Nurieva et al., 2009; Tangye et al., 2002). We also detected T_H_1 and CD8 T cells in BAL at week 8 that decreased to undetectable levels at week 39. Such responses were increased with either mRNA-1273 or mRNA-Omicron (Fig. S5).

### Boosting with mRNA-1273 or mRNA-Omicron provides equivalent protection in the lungs against Omicron challenge

To determine the extent of protection provided by a homologous mRNA-1273 or challenge virus-matched mRNA-Omicron boost following the two-dose mRNA-1273 immunization series, we obtained a new viral stock of Omicron, which was sequenced and confirmed to contain the canonical mutations present in the dominant Omicron sub-lineage BA.1 (Fig. S7).

Four weeks after administration of either boost, we challenged these NHP and 8 control NHPs with 1x10^6^ plaque forming units (PFU) via both intratracheal (IT) and intranasal (IN) routes. The control NHP had previously been administered 50µg of control mRNA formulated in lipid nanoparticles at the time of boost and had never been vaccinated.

BAL, nasal swabs (NS) and oral swabs (OS) were collected following challenge. Copy numbers of SARS-CoV-2 subgenomic RNA (sgRNA) were measured to determine the extent of viral replication. As sgRNA encoding for the N gene (sgRNA_N) are the most abundant transcripts produced due to the viral discontinuous transcription process (Kim et al., 2020), the sgRNA_N qRT-PCR assay was chosen for its enhanced sensitivity. On day 2 post-infection in the BAL, unvaccinated NHP had geometric mean copy numbers of 1x10^6^ sgRNA_N per mL whereas the vaccinated NHP had 3x10^2^ and 2x10^2^ for the mRNA-1273 and mRNA-Omicron cohorts, respectively (Fig. 5A). By day 4, all vaccinated NHP had undetectable levels of sgRNA_N while copy numbers in the unvaccinated group had only declined to 3x10^5^ per mL (mRNA-1273 vs control on days 2 and 4: *P*<0.0001; mRNA-Omicron vs control on days 2 and 4: *P*<0.0001).

**Figure 5.**
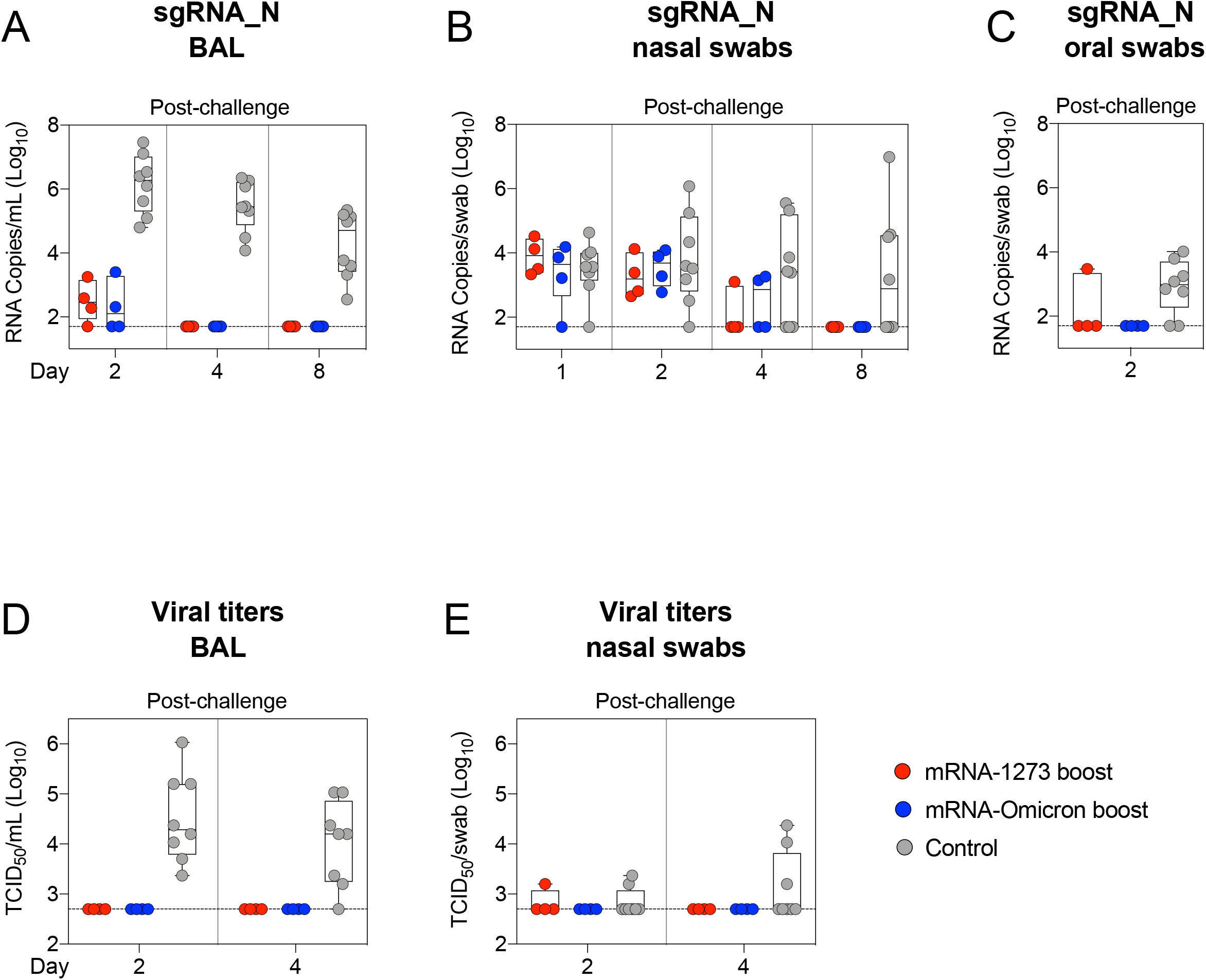
Boosting provides equivalent protection in the lungs against Omicron challenge. (A-E) BAL (A, D), NS (B, E) and OS (C) were collected at the indicated times following challenge with 1x10^6^ PFU Omicron. (A-C) Omicron sgRNA_N copy numbers per mL of BAL or per swab. (D-E) Viral titers per mL of BAL or per swab. Circles indicate individual NHP. Boxes represent interquartile range with the median denoted by a horizontal line. Assay LOD indicated by dotted lines. 8 controls and 4 vaccinated NHP per boost cohort. See also Figure S7 for Omicron challenge stock sequence.

In the nose, sgRNA_N copy numbers at days 1 to 4 were low for most animals and were not different between the control and vaccinated cohorts, so protection following vaccination could not be determined (Fig. 5B). At day 4, 5/8 controls had detectable virus in the nose as compared to 3/8 vaccinated NHP, with no clear difference between the boost cohorts. However, by day 8, 4/8 controls still had detectable sgRNA_N including 2 animals with increased copy numbers while none of the vaccinated NHP had detectable sgRNA.

In assessing sgRNA_N in the throat, it is noteworthy that 2 days after challenge, only 1/8 vaccinated NHP (in either boost group) had detectable virus in the throat compared to 6/8 control NHP (Fig. 5C).

We also measured the amount of culturable virus using a tissue culture infectious dose assay (TCID_50_). No virus was detected in the BAL of any vaccinated NHP, while 8/8 and 7/8 control NHP had detectable virus 2 and 4 days after challenge, respectively (Fig. 5D). In the NS, 1/8 boosted animals had culturable virus at any timepoint. In the unvaccinated control animals, 2/8 and 3/8 NHP had culturable virus in the nose 2 and 4 days after challenge, respectively (Fig. 5E).

### Viral antigen and pathology in the lungs after challenge

To assess lung pathology in NHP, 2 of the animals in each group were euthanized on day 8 following Omicron challenge, and the amount of viral antigen (SARS-CoV-2 N) and inflammation in the lungs were assessed (Fig. 6). N antigen was detected in variable amounts in the lungs of both control animals. When present, viral antigen was often associated with the alveolar capillaries and, occasionally, nearby immune cells. There was no evidence of viral antigen in the lungs of the vaccinated NHP.

**Figure 6.**
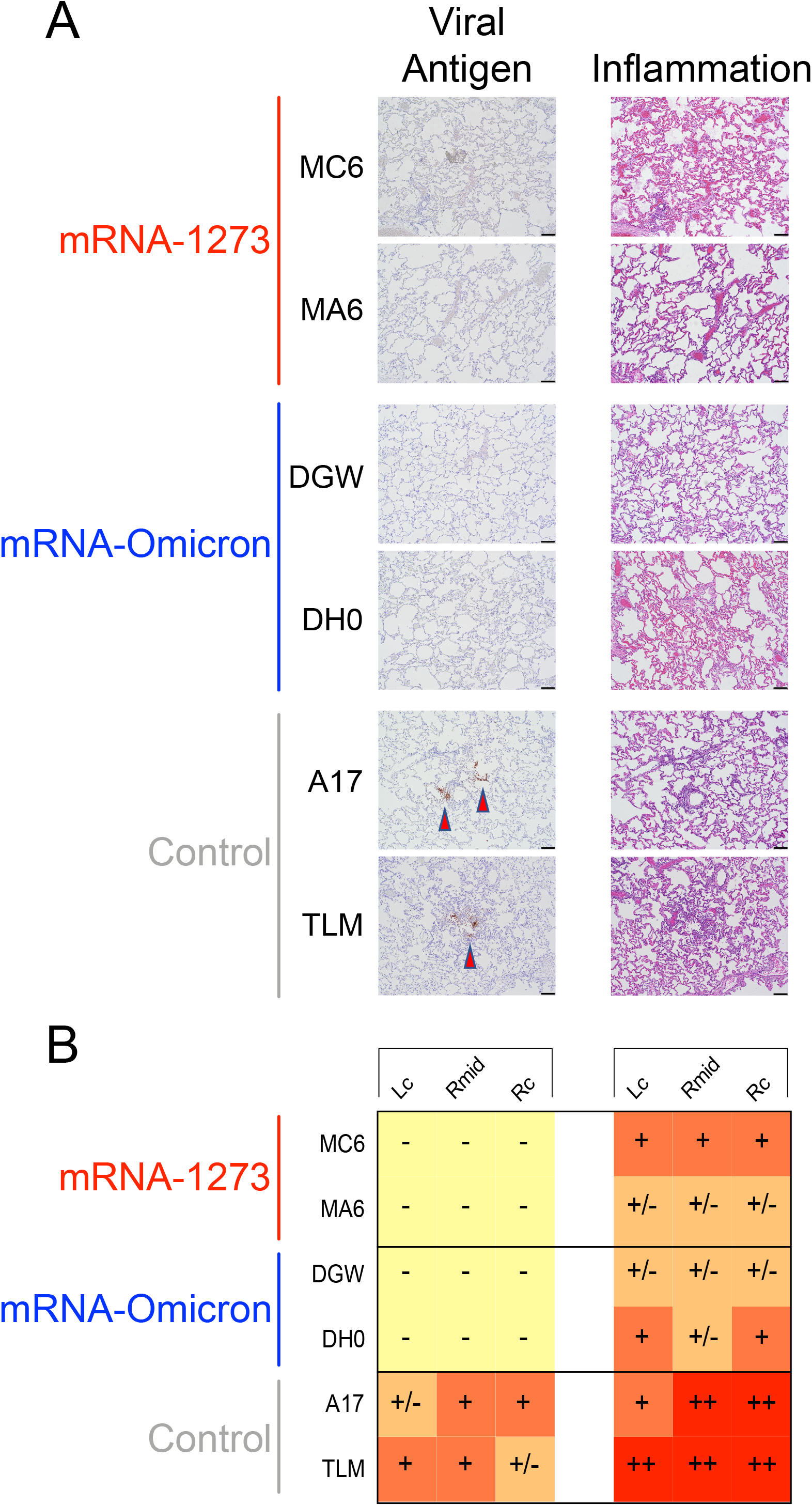
Viral antigen and pathology in the lungs after challenge. (A-B) 2 NHP per group were euthanized on day 8 post-challenge and tissue sections taken from lungs. (A) *Left.* Representative images indicating detection of SARS-CoV-2 N antigen by immunohistochemistry with a polyclonal anti-N antibody. Antigen-positive foci are marked by a red arrow. *Right.* Hematoxylin and eosin stain (H&E) illustrating the extent of inflammation and cellular infiltrates. Images at 10x magnification with black bars for scale (100µm). (B) SARS-CoV-2 antigen and inflammation scores in the left cranial lobe (Lc), right middle lobe (Rmid) and right caudal lobe (Rc) of the lungs. Antigen scoring legend: **-** no antigen detected; **+/-** rare to occasional foci; **+** occasional to multiple foci; **++** multiple to numerous foci; **+++** numerous foci. Inflammation scoring legend: - minimal to absent inflammation; **+/-** minimal to mild inflammation; **+** mild to moderate inflammation; **++** moderate to severe inflammation; **+++** severe inflammation. Horizontal rows correspond to individual NHP depicted above (A).

Animals from both boost groups displayed histopathologic alterations that were classified as minimal to mild or moderate. Inflammation was largely characterized by mild and patchy expansion of alveolar capillaries, generalized alveolar capillary hypercellularity, mild and regional type II pneumocyte hyperplasia and, less frequently, scattered collections of immune cells within some alveolar spaces. In contrast, unvaccinated animals were characterized as having a moderate to severe pathology. Lung sections from controls included features characterized by moderate and often diffuse alveolar capillary expansion, diffuse hypercellularity, moderate type II pneumocyte hyperplasia and multiple areas of perivascular cellular infiltration. Together, these data indicate that protection against Omicron was robust in the lungs regardless of boost selection.

## Discussion

Omicron has become the dominant global variant of SARS-CoV-2 due to its transmission advantage relative to Delta and its ability to evade prior immunity (Grabowski et al., 2022; Viana et al., 2022). Vaccine efficacy against infection with Omicron has declined and boosting with a third dose of an mRNA COVID-19 vaccine matched to the prototype strain has been shown to restore immunity and protection (Accorsi et al., 2022; Garcia-Beltran et al., 2022; Hansen et al., 2021; Pajon et al., 2022; Tseng et al., 2022). Here, we immunized NHP with 2 doses of mRNA-1273 (100μg) and boosted them ∼9 months later with 50μg of either mRNA-1273 or mRNA-Omicron prior to challenge with Omicron virus. The principal findings were: (1) 9 months after the two-dose regimen, neutralizing and binding antibody titers to Omicron had declined substantially in blood and mucosal airways; (2) after the boost, neutralizing antibody titers to ancestral strains were restored and those to Omicron were increased compared to the peak response after the initial prime and boost; (3) both boosts expanded cross-reactive memory B cells but only the homologous boost was capable of expanding B cells specific for epitopes unique to the ancestral strain; and (4) both boosts provided complete protection in the lungs and limited protection in the upper airway after Omicron challenge.

Following either mRNA-1273 or mRNA-Omicron boost, there was essentially complete protection in the lower airway with no culturable virus by day 2 and no detectable sgRNA_N by day 4. These data are comparable to our previous findings of equivalent upper and lower airway protection following Beta challenge in NHP boosted with either mRNA-1273 or mRNA-1273.Beta 1-2 months after immunization with 2 doses of mRNA-1273 (Corbett et al., 2021a). In contrast to the lower airway, there were no clear and consistent differences in sgRNA_N copy number at days 2 or 4 in the upper airway of vaccinated or control NHP. Of note, more of the control animals had detectable sgRNA at day 4 and increased sgRNA at day 8 as compared to the boosted animals. We would also note that the amount of Omicron replication as assessed by sgRNA or culturable virus in the control animals is demonstrably different than in our prior studies in which NHP were challenged with WA1, Delta or Beta (Corbett et al., 2020; Corbett et al., 2021c; Gagne et al., 2022). These findings compliment a growing body of evidence for reduced overall severity of Omicron infection in animal models of COVID-19 compared to other variants (Bentley et al., 2021; Halfmann et al., 2022; Suryawanshi et al., 2022). Overall, the findings here of high-level protection in the lungs recapitulate observations in mRNA-1273-vaccinated humans of reduced disease severity following infection with Omicron (Abdullah et al., 2021; Sigal, 2022; Wolter et al., 2022).

Neutralizing antibodies to Omicron in the blood or ACE2 binding inhibitory antibodies in the airway mucosa were low after the first 2 doses of mRNA-1273 at weeks 6-8 and low to undetectable ∼9 months later. Importantly, either mRNA-1273 or mRNA-Omicron boosts were able to significantly increase neutralizing antibody titers against Omicron and Beta beyond their initial peak consistent with a rapid recall B cell response. This also implies that neutralizing antibody titers at extended times after vaccination may not be a reliable surrogate either for vaccine efficacy in the lower airway or for predicting responses following a boost or infection as they may not reflect the recall capacity of the underlying memory B cell population.

The observation that boosting with either mRNA-1273 or mRNA-Omicron resulted in the expansion of a similarly high frequency of cross-reactive B cells likely stems from the principle of original antigenic sin, otherwise termed antigenic imprinting, whereby prior immune memory is recalled by a related antigenic encounter (Davenport and Hennessy, 1957; Davenport et al., 1953). Recall of prior immunity may be either deleterious or beneficial as exemplified by the impact of the circulating influenza A subtypes at the time of an individual’s first exposure after birth on patterns of disease susceptibility to subsequent pandemic influenza A outbreaks (Gostic et al., 2016; Worobey et al., 2014). The current worldwide distribution and evolution of SARS-CoV-2, however, is quite different from that of influenza A. Whereas multiple subtypes of influenza A circulate with different levels of co-dominance, SARS-CoV-2 distribution has generally become rapidly dominated by a single variant — currently Omicron — before replacement by another that, for various reasons, is more transmissible. The question therefore is whether there is added value from boosting with a heterologous vaccine matched to the dominant circulating variant, or whether cross-reactive B cell recall immunity elicited by boosting with the original vaccine is sufficient to reduce infection and disease severity. As we have now shown in two different NHP studies, boosting animals with either mRNA-1273.Beta (Corbett et al., 2021a) or mRNA-Omicron provided no advantage over mRNA-1273 in recalling high titer neutralizing antibodies across all variants tested and protecting from virus replication after challenge. These considerations apply to the large numbers of individuals with prior immunity from vaccination or infection with current and previous variants who may not necessarily benefit from a change in vaccine design.

Looking to the future, however, if Omicron, or a closely antigenically related variant, remains the dominant circulating variant for some years to come, then it is possible that a change in the initial vaccine regimen would be warranted, particularly in immunologically naïve populations such as children as they reach the age of eligibility for approved COVID-19 vaccines. Importantly, it would need to be established that a switch in COVID-19 vaccine design to match the current dominant variant would not jeopardize responses against variants which may be antigenically distant from Omicron but close to the prototype. Indeed a recent study has shown that immunization of mice with an Omicron mRNA vaccine elicited strong neutralization against Omicron but not to other variants, consistent with data from primary Omicron infection (Lee et al., 2022; Roessler et al., 2022; Suryawanshi et al., 2022). Thus, a combination or bivalent vaccine to generate B cells specific for the current variant as well as cross-reactive to other variants might ensure greater breadth of neutralization.

In summary, our findings highlight two important factors that will impact management of this pandemic. The first is the design of the vaccine and whether it should be changed based on the currently circulating variant. At present, boosting with mRNA-1273 provides robust increases in neutralizing antibodies and appears to be sufficient to prevent severe disease after exposure from all known variants (Baden et al., 2021a; Bruxvoort et al., 2021a; Bruxvoort et al., 2021b; Chemaitelly et al., 2021; Corbett et al., 2020; Corbett et al., 2021c; Gagne et al., 2022; Pilishvili et al., 2021; Tang et al., 2021). Variant-matched vaccines may be preferable in the future if new variants were to emerge that were even further antigenically distant such that cross-reactive epitopes are rendered ineffective or if there were differences in the durability of neutralizing antibody titers elicited by different boosts. Second, as neutralizing antibody titers wane with time after vaccination, their ability to serve as a surrogate for vaccine efficacy or to predict clinical outcomes against severe disease after infection with VOC may become diminished. Thus the determination of when to administer a boost may depend on the recall capacity of the underlying memory B cell population. These considerations will become clear as human clinical data are made available.

### Limitations of the study

There are several limitations to this study. First, NHP models may not fully recapitulate clinical data in humans regarding the extent of viral replication necessary for the enhanced transmission of Omicron compared to prior variants. Even if a significant component of Omicron’s growth advantage is due to immune escape, the role of viral replication kinetics cannot be ruled out.

Here, viral titers were low in the lungs and low to undetectable in the upper airway. Second, neutralizing antibody titers in NHP are 5-to 10-fold greater than in humans that received the same dose and regimen of mRNA-1273 with a boost (Edara et al., 2021a; Pajon et al., 2022). Third, a second dose of mRNA-Omicron may elicit a population of B cells specific only for Omicron. Finally, since we sought to compare two different mRNA boosts, we did not have an unboosted group to determine whether the boost enhanced protection. As all the boosted NHP were completely protected in the lungs, we were unable to determine an immune threshold for protection.

## Materials and methods

### Resource availability

#### Lead contact

Further information and requests for resources should be directed to and will be fulfilled by the lead contact, Robert A. Seder.

#### Materials availability

This study did not generate new unique reagents.

#### Data and code availability

• All data reported in this paper will be shared by the lead contact upon request.
• This paper does not report original code.
• Any additional information required to reanalyze the data reported in this paper is available from the lead contact upon request.

### Experimental model and subject details

#### Preclinical mRNA and lipid nanoparticle production

A sequence-optimized mRNA encoding prefusion-stabilized SARS-CoV-2 S protein containing 2 proline stabilization mutations (S-2P) (Pallesen et al., 2017; Wrapp et al., 2020) for WA1 and Omicron were synthesized in vitro and formulated (Hassett et al., 2019). Control mRNA “UNFIX-01 (Untranslated Factor 9)” was synthesized and similarly formulated into lipid nanoparticles as previously described (Corbett et al., 2021a).

#### Rhesus macaque model and immunizations

All experiments conducted according to NIH regulations and standards on the humane care and use of laboratory animals as well as the Animal Care and Use Committees of the NIH Vaccine Research Center and BIOQUAL, Inc. (Rockville, Maryland). All studies were conducted at BIOQUAL, Inc. Four- to eight-year-old rhesus macaques of Indian origin were stratified into groups based on sex, age and weight. Eight macaques were immunized with mRNA-1273 at weeks 0 and 4 with a dose of 100μg delivered intramuscularly in 1mL of formulated lipid nanoparticles diluted in phosphate-buffered saline (PBS) into the right quadricep as previously described (Corbett et al., 2020; Corbett et al., 2021c; Gagne et al., 2022). At week 41 (∼9 months after the second immunization), the eight macaques were split into two groups of 4 and boosted with 50μg mRNA-1273 or 50μg of mRNA-Omicron. Animals in the control group were immunized with 50μg control mRNA at the time of the boost.

### Method details

#### Cells and viruses

VeroE6-TMPRSS2 cells were generated at the Vaccine Research Center, NIH, Bethesda, MD. Isolation and sequencing of EHC-083E (D614G SARS-CoV-2), Delta, Beta and Omicron for live virus neutralization assays were previously described (Edara et al., 2021b; Edara et al., 2021c; Vanderheiden et al., 2020). Viruses were propagated in Vero-TMPRSS2 cells to generate viral stocks. Viral titers were determined by focus-forming assay on VeroE6-TMPRSS2 cells. Viral stocks were stored at -80°C until use.

#### Sequencing of Omicron virus stock

We used NEBNext Ultra II RNA Prep reagents and multiplex oligos (New England Biolabs) to prepare Illumina-ready libraries, which were sequenced on a MiSeq (Illumina) as described previously (Corbett et al., 2021c; Gagne et al., 2022). Demultiplexed sequence reads were analyzed in the CLC Genomics Workbench v.21.0.3 by (1) trimming for quality, length, and adaptor sequence, (2) mapping to the Wuhan-Hu-1 SARS-CoV-2 reference (GenBank no. NC_045512), (3) improving the mapping by local realignment in areas containing insertions and deletions (indels), and (4) generating both a sample consensus sequence and a list of variants. Default settings were used for all tools.

#### Omicron challenge

Macaques were challenged at week 45 (4 weeks after the second boost) with a total dose of 1 × 10^6^ PFU of SARS-CoV-2 Omicron. The viral inoculum was administered as 7.5 ×10^5^ PFUs in 3mL intratracheally and 2.5 ×10^5^ PFUs in 1mL intranasally in a volume of 0.5mL distributed evenly into each nostril.

#### Serum and mucosal antibody titers

Quantification of antibodies in the blood and mucosa were performed as previously described (Corbett et al., 2020). Briefly, total IgG antigen-specific antibodies to variant SARS-CoV-2 S- and RBD-derived antigens were determined in a multiplex serology assay by MSD V-Plex SARS-CoV-2 Panel 23 for S and MSD V-Plex SARS-CoV-2 Panel 22 for RBD) according to manufacturer’s instructions, except 25μl of sample and detection antibody were used per well. Heat inactivated plasma was initially diluted 1:100 and then serially diluted 1:10 for blood S- and 1:4 for RBD-binding. BAL fluid and nasal washes were concentrated 10-fold with Amicon Ultra centrifugal filter devices (Millipore Sigma). Concentrated samples were diluted 1:5 prior to 5-fold serial dilutions.

#### S-2P antigens

While S antigens were used for binding ELISAs, S-2P antigens were used for ACE2 inhibition assays and B cell probe binding. S-2P constructs were made as follows. Biotinylated S probes were expressed transiently for WA1, Delta, Beta and Omicron strains and purified and biotinylated in a single in-process step (Teng et al., 2021; Zhou et al., 2020). S-2P for WA1 and Omicron were made as previously described (Olia et al., 2021).

#### S-2P-ACE2 binding inhibition

ACE2 binding inhibition was performed using a modified Meso Scale Discovery (MSD) platform assay. Briefly, after blocking MSD Streptavidin MULTI-ARRAY 384 well plates with Blocker A (MSD), the plates were coated with 1 µg/ml of biotinylated SARS-CoV-2 variant S-2P (WA1, Beta, Delta or Omicron) and incubated for 1 hour at room temperature (RT). The plates were washed 5 times with wash buffer (1x PBS containing 0.05% Tween-20). Diluted samples were added to the coated plates and incubated for 1 hour at RT. MSD SULFO-TAG Human ACE2 protein was diluted 1:200 and added to the plates. After 1 hour incubation at RT, the plates were washed 5 times with wash buffer and read on MSD Sector S 600 instrument after the addition of Gold Read Buffer B (MSD). Results are reported as percent inhibition. BAL fluid and nasal washes were first concentrated 10-fold with Amicon Ultra centrifugal filter devices (Millipore Sigma) and then diluted 1:5 in Diluent 100 (MSD).

#### Focus reduction neutralization assay

FRNT assays were performed as previously described (Edara et al., 2021b; Edara et al., 2021c; Vanderheiden et al., 2020). Briefly, samples were diluted at 3-fold in 8 serial dilutions using DMEM (VWR, #45000-304) in duplicates with an initial dilution of 1:10 in a total volume of 60μl. Serially diluted samples were incubated with an equal volume of WA1, Delta, Beta or Omicron (100-200 foci per well based on the target cell) at 37°C for 45 minutes in a round-bottomed 96-well culture plate. The antibody-virus mixture was then added to VeroE6-TMPRSS2 cells and incubated at 37°C for 1 hour. Post-incubation, the antibody-virus mixture was removed and 100µl of pre-warmed 0.85% methylcellulose overlay was added to each well. Plates were incubated at 37°C for 18 hours and the methylcellulose overlay was removed and washed six times with PBS. Cells were fixed with 2% paraformaldehyde in PBS for 30 minutes. Following fixation, plates were washed twice with PBS and permeabilization buffer (0.1% BSA, 0.1% Saponin in PBS) was added to permeabilized cells for at least 20 minutes. Cells were incubated with an anti-SARS-CoV S primary antibody directly conjugated to Alexaflour-647 (CR3022-AF647) overnight at 4°C. Foci were visualized and imaged on an ELISPOT reader (CTL). Antibody neutralization was quantified by counting the number of foci for each sample using the Viridot program (Katzelnick et al., 2018). The neutralization titers were calculated as follows: 1 - (ratio of the mean number of foci in the presence of sera and foci at the highest dilution of respective sera sample). Each specimen was tested in duplicate. The FRNT-50 titers were interpolated using a 4-parameter nonlinear regression in GraphPad Prism 9.2.0. Samples that do not neutralize at the limit of detection (LOD) at 50% are plotted at 20 and was used for geometric mean and fold-change calculations. The assay LOD was 20.

#### Lentiviral pseudovirus neutralization

Neutralizing antibodies in serum or plasma were measured in a validated pseudovirus-based assay as a function of reductions in luciferase reporter gene expression after a single round of infection with SARS-CoV-2 spike-pseudotyped viruses in 293T/ACE2 cells (293T cell line stably overexpressing the human ACE2 cell surface receptor protein, obtained from Drs. Mike Farzan and Huihui Mu at Scripps) as previously described (Gilbert et al., 2021; Shen et al., 2021). SARS-CoV-2 S-pseudotyped virus was prepared by transfection in 293T/17 cells (human embryonic kidney cells in origin; obtained from American Type Culture Collection, #CRL-11268) using a lentivirus backbone vector, a spike-expression plasmid encoding S protein from Wuhan-Hu-1 strain (GenBank no. NC_045512) with a p.Asp614Gly mutation, a TMPRSS2 expression plasmid and a firefly Luc reporter plasmid. For pseudovirus encoding the S from Delta, Beta and Omicron, the plasmid was altered via site-directed mutagenesis to match the S sequence to the corresponding variant sequence as previously described (Corbett et al., 2021a). A pre-titrated dose of pseudovirus was incubated with eight serial 5-fold dilutions of serum samples (1:20 start dilution) in duplicate in 384-well flat-bottom tissue culture plates (Thermo Fisher, #12-565-344) for 1 hour at 37°C prior to adding 293T/ACE2 cells. One set of 14 wells received cells and virus (virus control) and another set of 14 wells received cells only (background control), corresponding to technical replicates. Luminescence was measured after 66-72 hours of incubation using Britelite-Plus luciferase reagent (Perkin Elmer, #6066769). Neutralization titers are the inhibitory dilution of serum samples at which relative luminescence units (RLUs) were reduced by 50% (ID_50_) compared to virus control wells after subtraction of background RLUs. Serum samples were heat-inactivated for 30-45 minutes at 56°C prior to assay.

#### Serum antibody avidity

Avidity was measured as described previously (Francica et al., 2021) in an adapted ELISA assay. Briefly, ELISA against S-2P was performed in the absence or presence of sodium thiocyanate (NaSCN) and developed with HRP-conjugated goat anti-monkey IgG (H+L) secondary antibody (Invitrogen) and SureBlue 3,3′,5,5′-tetramethylbenzidine (TMB) microwell peroxidase substrate (1-Component; SeraCare) and quenched with 1 N H_2_SO_4_. The avidity index (AI) was calculated as the ratio of IgG binding to S-2P in the absence or presence of NaSCN.

#### Epitope mapping

Serum epitope mapping competition assays were performed, as previously described (Corbett et al., 2021a), using the Biacore 8K+ surface plasmon resonance system (Cytiva). Briefly, through primary amine coupling using a His capture kit (Cytiva), anti-histidine antibody was immobilized on Series S Sensor Chip CM5 (Cytiva) allowing for the capture of his-tagged SARS-CoV-2 S-2P on active sensor surface.

Human IgG monoclonal antibodies (mAbs) used for these analyses include: RBD-specific mAbs B1-182, A19-46.1, A20-29.1, A19-61.1, S309, A23-97.1 and A23-80.1. Negative control antibody or competitor mAb was injected over both active and reference surfaces. Following this, NHP sera (diluted 1:50) was flowed over both active and reference sensor surfaces. Active and reference sensor surfaces were regenerated between each analysis cycle.

For analysis, sensorgrams were aligned to Y (Response Units) = 0, using Biacore 8K Insights Evaluation Software (Cytiva) beginning at the serum association phase. Relative “analyte binding late” report points (RU) were collected and used to calculate fractional competition (% C) using the following formula: % C = [1 – (100 * ( (RU in presence of competitor mAb) / (RU in presence of negative control mAb) ))]. Results are reported as fractional competition. Assays were performed in duplicate, with average data point represented on corresponding graphs.

#### B cell probe binding

Kinetics of S-specific memory B cells responses were determined as previously described (Gagne et al., 2022). Briefly, cryopreserved PBMC were thawed and stained with the following antibodies (monoclonal unless indicated): IgD FITC (goat polyclonal, Southern Biotech), IgM PerCP-Cy5.5 (clone G20-127, BD Biosciences), IgA Dylight 405 (goat polyclonal, Jackson Immunoresearch Inc), CD20 BV570 (clone 2H7, Biolegend), CD27 BV650 (clone O323, Biolegend), CD14 BV785 (clone M5E2, Biolegend), CD16 BUV496 (clone 3G8, BD Biosciences), CD4 BUV737 (clone SK3, BD Biosciences), CD19 APC (clone J3-119, Beckman), IgG Alexa 700 (clone G18-145, BD Biosciences), CD3 APC-Cy7 (clone SP34-2, BD Biosciences), CD38 PE (clone OKT10, Caprico Biotechnologies), CD21 PE-Cy5 (clone B-ly4, BD Biosciences) and CXCR5 PE-Cy7 (clone MU5UBEE, Thermo Fisher Scientific). Stained cells were then incubated with streptavidin-BV605 (BD Biosciences) labeled WA1, Beta, Delta or Omicron S-2P and streptavidin-BUV661 (BD Biosciences) labeled WA1 or Delta S-2P for 30 minutes at 4°C (protected from light). Cells were washed and fixed in 0.5% formaldehyde (Tousimis Research Corp) prior to data acquisition. Aqua live/dead fixable dead cell stain kit (Thermo Fisher Scientific) was used to exclude dead cells. All antibodies were previously titrated to determine the optimal concentration. Samples were acquired on an BD FACSymphony cytometer and analyzed using FlowJo version 10.7.2 (BD, Ashland, OR).

#### Intracellular cytokine staining

Intracellular cytokine staining was performed as previously described (Donaldson et al., 2019; Gagne et al., 2022). Briefly, cryopreserved PBMC and BAL cells were thawed and rested overnight in a 37°C/5% CO_2_ incubator. The following morning, cells were stimulated with SARS-CoV-2 S protein (S1 and S2, matched to vaccine insert) peptide pools (JPT Peptides) at a final concentration of 2 μg/ml in the presence of 3mM monensin for 6 hours. The S1 and S2 peptide pools were comprised of 158 and 157 individual peptides, respectively, as 15 mers overlapping by 11 amino acids in 100% DMSO. Negative controls received an equal concentration of DMSO instead of peptides (final concentration of 0.5%). The following monoclonal antibodies were used: CD3 APC-Cy7 (clone SP34-2, BD Biosciences), CD4 PE-Cy5.5 (clone S3.5, Invitrogen), CD8 BV570 (clone RPA-T8, BioLegend), CD45RA PE-Cy5 (clone 5H9, BD Biosciences), CCR7 BV650 (clone G043H7, BioLegend), CXCR5 PE (clone MU5UBEE, Thermo Fisher), CXCR3 BV711 (clone 1C6/CXCR3, BD Biosciences), PD-1 BUV737 (clone EH12.1, BD Biosciences), ICOS Pe-Cy7 (clone C398.4A, BioLegend), CD69 ECD (cloneTP1.55.3, Beckman Coulter), IFN-g Ax700 (clone B27, BioLegend), IL-2 BV750 (clone MQ1-17H12, BD Biosciences), IL-4 BB700 (clone MP4-25D2, BD Biosciences), TNF-FITC (clone Mab11, BD Biosciences), IL-13 BV421 (clone JES10-5A2, BD Biosciences), IL-17 BV605 (clone BL168, BioLegend), IL-21 Ax647 (clone 3A3-N2.1, BD Biosciences) and CD154 BV785 (clone 24-31, BioLegend). Aqua live/dead fixable dead cell stain kit (Thermo Fisher Scientific) was used to exclude dead cells. All antibodies were previously titrated to determine the optimal concentration. Samples were acquired on a BD FACSymphony flow cytometer and analyzed using FlowJo version 10.8.0 (BD, Ashland, OR).

#### Subgenomic RNA quantification

sgRNA was isolated and quantified by researchers blinded to vaccine history as previously described (Corbett et al., 2021c), except for the use of a new probe noted below. Briefly, total RNA was extracted from BAL fluid and nasal swabs using RNAzol BD column kit (Molecular Research Center). PCR reactions were conducted with TaqMan Fast Virus 1-Step Master Mix (Applied Biosystems), forward primer in the 5’ leader region and gene-specific probes and reverse primers as follows:

sgLeadSARSCoV2_F: 5’-CGATCTCTTGTAGATCTGTTCTC-3’

*N gene*

N2_P: 5’-FAM-CGATCAAAACAACGTCGGCCCC-BHQ1-3’ wtN_R: 5’-GGTGAACCAAGACGCAGTAT-3’

Amplifications were performed with a QuantStudio 6 Pro Real-Time PCR System (Applied Biosystems). The assay lower LOD was 50 copies per reaction.

#### Median Tissue Culture Infectious Dose (TCID_50_) assay

TCID_50_ was quantified as previously described (Corbett et al., 2021c). Briefly, Vero-TMPRSS2 cells were plated and incubated overnight. The following day, BAL samples were serially diluted, and the plates were incubated at 37 °C/5.0% CO2 for four days. Positive (virus stock of known infectious titer in the assay) and negative (medium only) control wells were included in each assay setup. The cell monolayers were visually inspected for cytopathic effect. TCID_50_ values were calculated using the Reed–Muench formula.

#### Histopathology and immunohistochemistry

Routine histopathology and detection of SARS-CoV-2 virus antigen via immunohistochemistry (IHC) were performed as previously described (Corbett et al., 2020; Gagne et al., 2022). Briefly, seven to nine days following Omicron challenge animals were euthanized and lung tissue was processed and stained with hematoxylin and eosin for pathological analysis or with a rabbit polyclonal anti-SARS-CoV-2 anti-nucleocapsid antibody (GeneTex, GTX135357) at a dilution of 1:2000 for IHC. Tissue sections were analyzed by a blinded board-certified veterinary pathologist using an Olympus BX43 light microscope. Photomicrographs were taken on an Olympus DP27 camera.

### Quantification and statistical analysis

Comparisons between groups, or between timepoints within a group, are based on unpaired and paired t-tests, respectively. All analysis for serum epitope mapping was performed using unpaired, two-tailed t-test. Binding, neutralizing and viral assays are log-transformed as appropriate and reported with geometric means and corresponding confidence intervals where indicated. There are no adjustments for multiple comparisons, so all p-values should be interpreted as suggestive rather than conclusive. All analyses are conducted using R version 4.0.2 and GraphPad Prism version 8.2.0 unless otherwise specified.

*P* values are stated in the text, and the sample *n* is listed in corresponding figure legends. For all data presented, *n*=4 for individual boost cohorts and *n*=7-8 for controls and vaccinated NHP at pre-boost timepoints.

## Acknowledgments

We would like to thank G. Alvarado for experimental organization and administrative support. The VRC Production Program (VPP) provided the WA1 protein for the avidity assay. VPP contributors include C. Anderson, V. Bhagat, J. Burd, J. Cai, K. Carlton, W. Chuenchor, N. Clbelli, G. Dobrescu, M. Figur, J. Gall, H. Geng, D. Gowetski, K. Gulla, L. Hogan, V. Ivleva, S. Khayat, P. Lei, Y. Li, I. Loukinov, M. Mai, S. Nugent, M. Pratt, E. Reilly, E. Rosales-Zavala, E. Scheideman, A. Shaddeau, A. Thomas, S. Upadhyay, K. Vickery, A. Vinitsky, C. Wang, C. Webber and Y. Yang.

This project has been funded in part by both the Intramural Program of the National Institute of Allergy and Infectious Diseases, National Institutes of Health, Department of Health and Human Services and under HHSN272201400004C (NIAID Centers of Excellence for Influenza Research and Surveillance, CEIRS) and NIH P51 OD011132 awarded to Emory University. This work was also supported in part by the Emory Executive Vice President for Health Affairs Synergy Fund award, COVID-Catalyst-I3 Funds from the Woodruff Health Sciences Center and Emory School of Medicine, the Pediatric Research Alliance Center for Childhood Infections and Vaccines and Children’s Healthcare of Atlanta, and Woodruff Health Sciences Center 2020 COVID-19 CURE Award.

## Author contributions

M.R., N.J.S., D.C.D. and R.A.S. designed experiments. M.G., J.I.M, K.E.F., S.F.A., B.J.F., A.P.W, D.A.W., B.C.L., C.M., N.J-B., R.C., S.L.F., M.P., M.E., V-V.E., N.V.M., M.M., L.M., C.C.H., B.M.N., K.W.B., C.N.M.D., J.C., J-P.M.T., E.M., L.P., A.V.R., B.N., D.V., A.C., A.D., K.S., D.R.F., S.T.N., S.G., A.R.H., F.L., J.R-T., C.G.L, S.A., D.K.E., H.A., M.G.L., K.S.C., M.C.N., A.B.M., M.S.S., I.N.M., M.R., N.J.S., D.C.D. and R.A.S. performed, analyzed, and/or supervised experiments. M.G., J.I.M., K.E.F., S.F.A., D.A.W., I.N.M. and D.C.D. designed figures. I-T.T., J.T., M.N., M.B., J.W., L.W., W.S., N.A.D-R., A.S.O., C.L., D.R.H., A.C., J.R.M. and P.D.K. provided critical reagents. M.G., J.I.M., D.C.D. and R.A.S. wrote manuscript. All authors edited the manuscript and provided feedback on research.

