## Supplemental Data for "mRNA-1273 or mRNA-Omicron boost in vaccinated macaques elicits comparable B cell expansion, neutralizing antibodies and protection against Omicron"

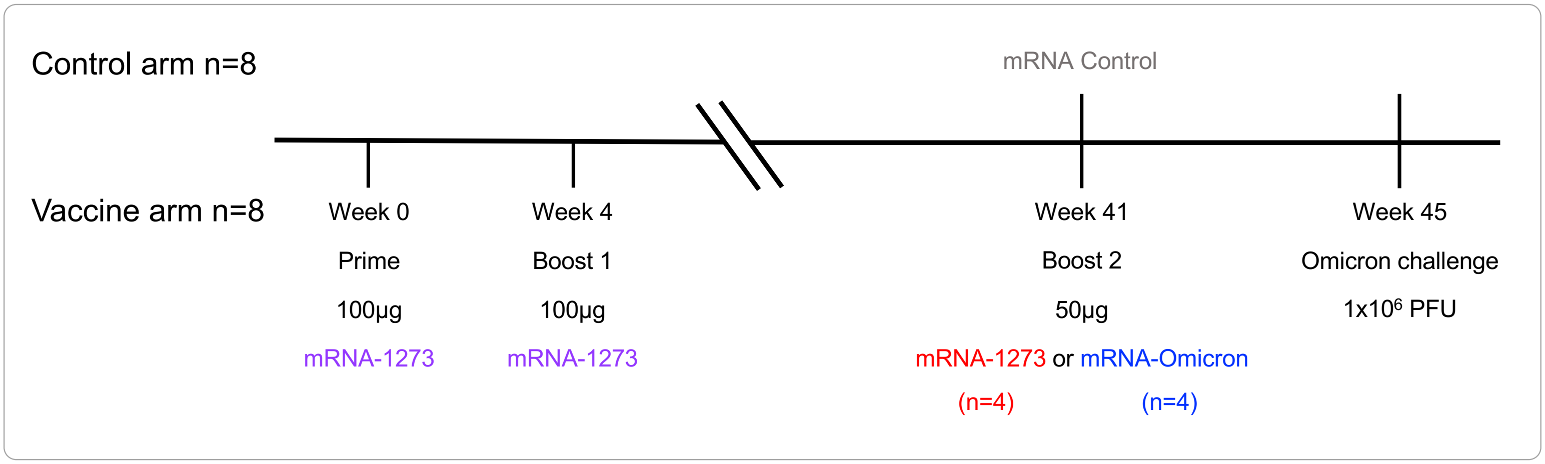


**Figure S1 Experimental timeline, related to Figure 1**

8 NHP were vaccinated with 100µg of mRNA-1273 at weeks 0 and 4. At week 41, NHP were split into 2 groups of 4 and boosted with 50µg of mRNA-1273 or mRNA-Omicron. Both groups, and 8 unvaccinated NHP which were given 50µg of mRNA control, were challenged with Omicron 1 month later.

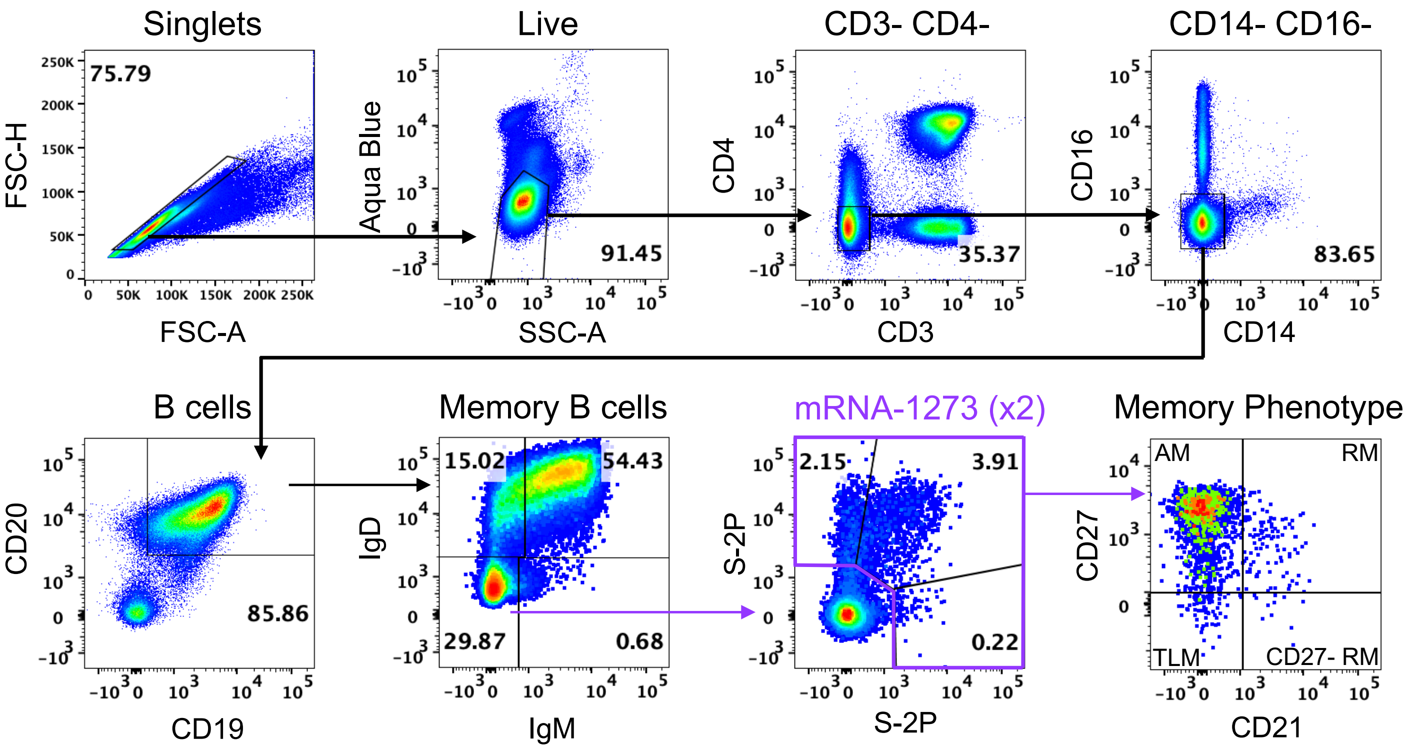


**Figure S2 B cell gating strategy, related to Figure 3**

Representative flow cytometry plots showing gating strategy for B cells in Figures 3 and 4 and Figure S3. Cells were gated as singlets and live cells on forward and side scatter and a live/dead aqua blue stain. CD3-, CD4- cells were then gated on absence of CD14 and CD16 expression and positive expression of CD20 and CD19. Memory B cells were selected based on lack of IgD or IgM. Finally, pairs of variant S-2P probes were used to determine binding specificity. Probe-binding cells were further characterized as having a phenotype consistent with CD27-negative resting memory (CD27- RM), tissue-like memory (TLM), activated memory (AM) or resting memory (RM) cells according to expression of CD27 and CD21.

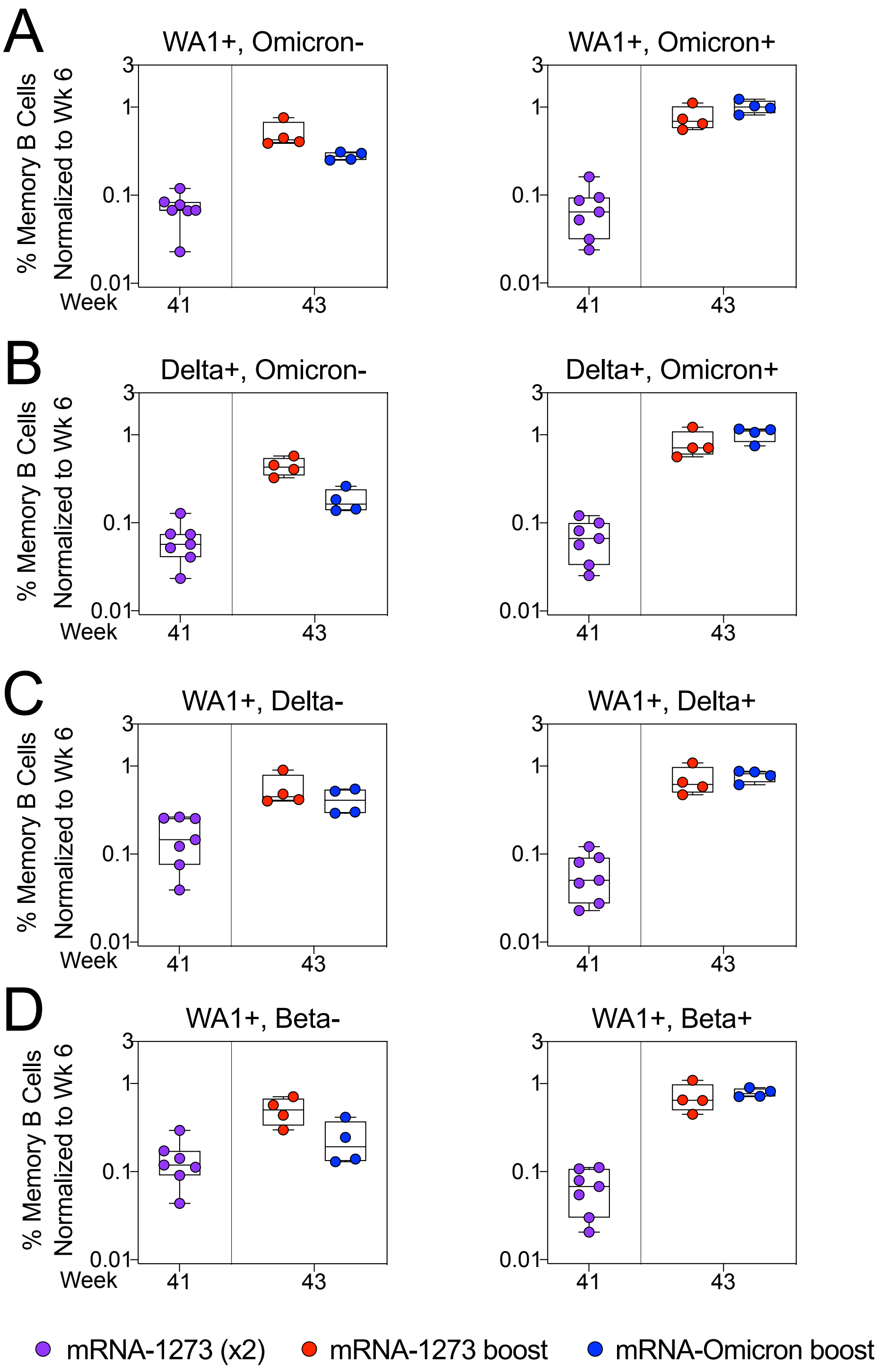
A
WA1+, Omicron-
WA1+, Omicron+
B
Delta+, Omicron-
Delta+, Omicron+
C
WA1+, Delta-
WA1+, Delta+
D
WA1+, Beta-
WA1+, Beta+
% Memory B Cells Normalized to Wk 6
3
1
0.1
0.01
Week
41
43
mRNA-1273 (x2)
mRNA-1273 boost
mRNA-Omicron boost

**Figure S3**

**Figure S3 Expansion of memory B cells that recognize unique WA1 epitopes only occurs after homologous mRNA-1273 boosting, related to Figure 4**

(A-D) Frequencies of memory B cells with indicated specificities as a percentage of total class-switched memory B cells (both S-2P-binding and non-S-2P-binding) were normalized to the corresponding frequencies from each individual NHP at week 6 post-immunization. Cross-reactivity shown for (A) WA1 and Omicron S-2P, (B) Delta and Omicron S-2P, (C) WA1 and Delta S-2P and (D) WA1 and Beta S-2P. Specificities not shown if memory B cell populations were indistinguishable from background staining. Circles indicate individual NHP. Boxes represent interquartile range with the median denoted by a horizontal line. A frequency of 1 indicates parity. 4-7 NHP per group.

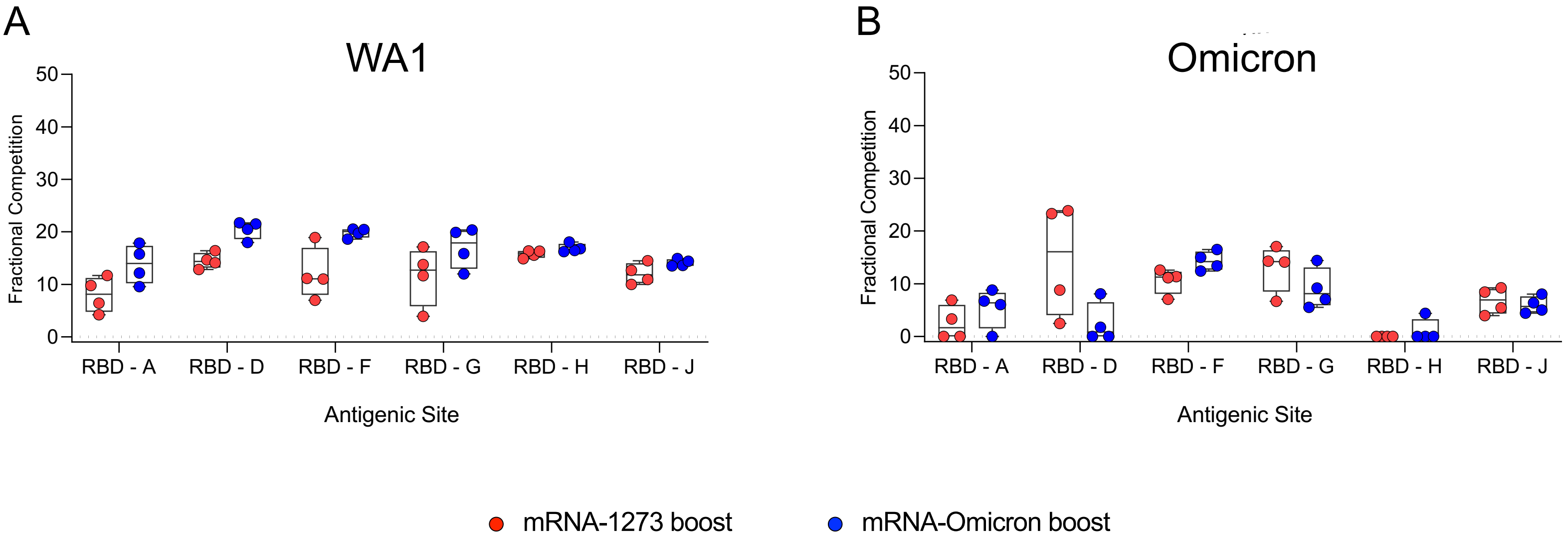


**Figure S4 Serum epitope reactivity following boost, related to Figure 4**

(A-B) Relative serum reactivity was measured as fractional competition of total measured serum antibody S-2P binding competed by single monoclonal antibody (mAb) targeting epitopes on WA1 (A) or Omicron (B) S-2P at week 43 post-immunization. Antigenic sites are defined by mAbs B1-182 (RBD-A), A19-46.1 (RBD-D), A19-61.1 (RBD-F), S309 (RBD-G), A23-97.1 (RBD-H) and A23-80.1 (RBD-J). Circles indicate individual NHP. Boxes represent interquartile range with the median denoted by a horizontal line. Dotted lines are for visualization purposes and denote 0% competition. 4 NHP per boost group.

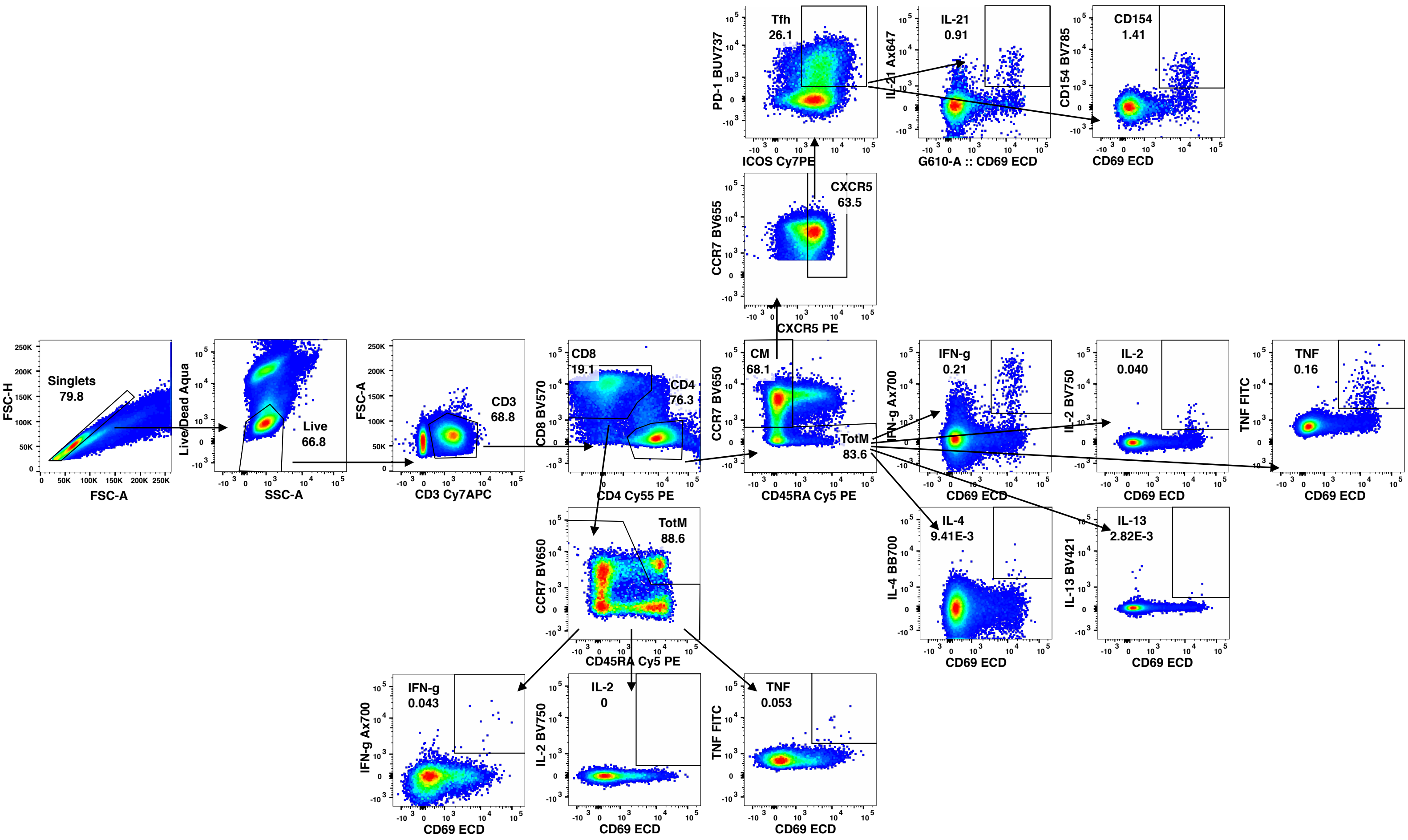


**Figure S5 T cell gating strategy, related to Figure 4**

Representative flow cytometry plots showing gating strategy for T cells in Figure S6. Cells were gated as singlets and live cells on forward and side scatter and a live/dead aqua blue stain. CD3$^{+}$ events were gated as CD4$^{+}$ or CD8$^{+}$ T cells. Total memory CD8$^{+}$ T cells were selected based on expression of CCR7 and CD45RA. Finally, SARS-CoV-2 S-specific memory CD8$^{+}$ T cells were gated according to co-expression of CD69 and IL-2, TNF or IFNγ. The CD4$^{+}$ events were defined as naïve, total memory or central memory according to expression of CCR7 and CD45RA. CD4$^{+}$ cells with a $T_H1$ phenotype were defined as memory cells that co-expressed CD69 and IL-2, TNF or IFNγ. CD4$^{+}$ cells with a $T_H2$ phenotype were defined as memory cells that co-expressed CD69 and IL-4 or IL-13. $T_{FH}$ cells were defined as central memory CD4$^{+}$ T cells that expressed CXCR5, ICOS and PD-1. $T_{FH}$ cells were further characterized as IL-21$^{+}$, CD69$^{+}$ or CD40L$^{+}$, CD69$^{+}$.

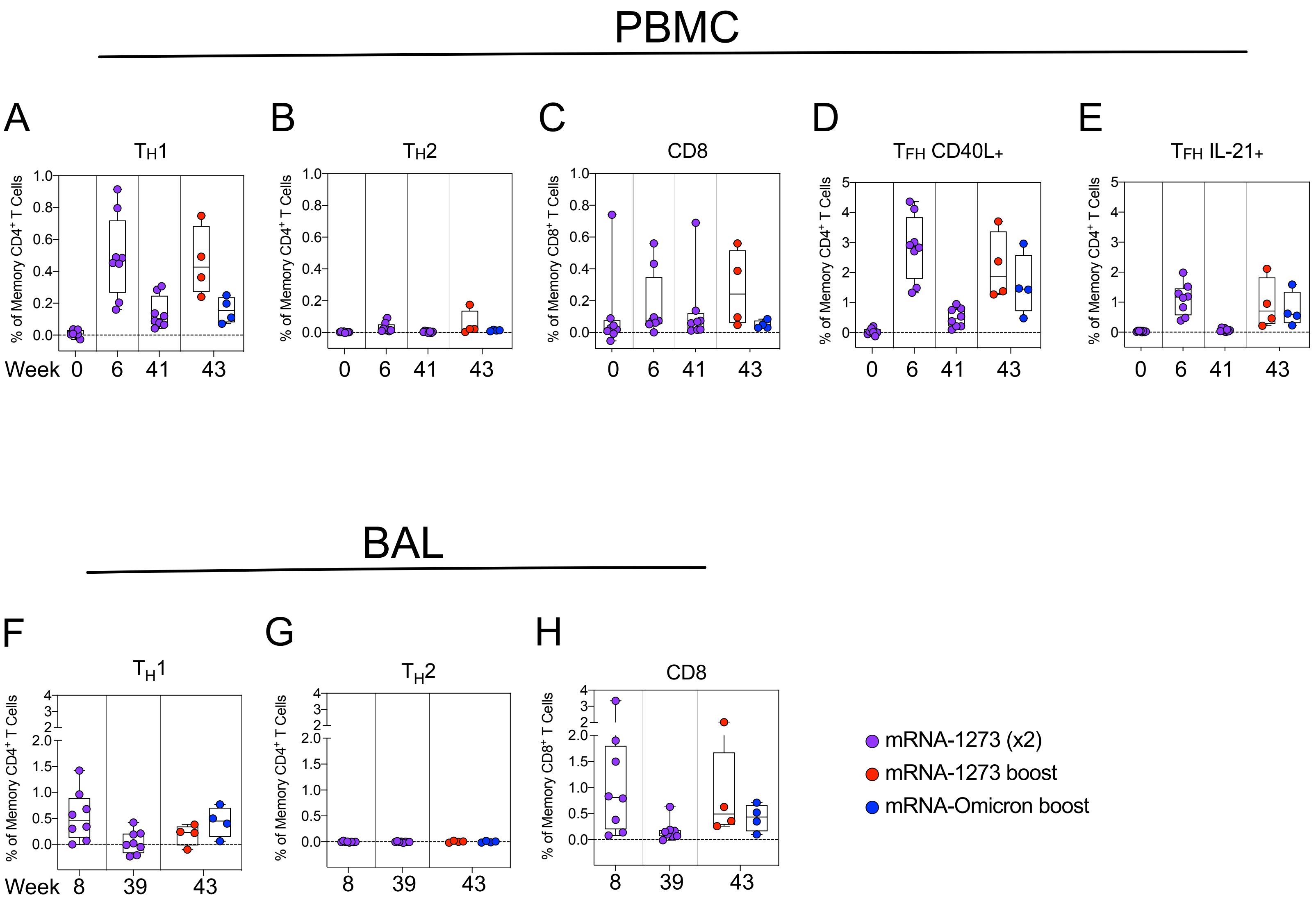

PBMC
A
B
C
D
E
TH1
TH2
CD8
TFH CD40L+
TFH IL-21+
% of Memory CD4+ T Cells
% of Memory CD8+ T Cells
Week 0 6 41 43
BAL
F
G
H
TH1
TH2
CD8
Week 8 39 43
mRNA-1273 (x2)
mRNA-1273 boost
mRNA-Omicron boost


Figure S6

**Figure S6 Both mRNA-1273 and mRNA-Omicron boost T cell responses to S peptides, related to Figure 4**

(A-E) PBMC collected at weeks 0, 6, 41 and 43 post-immunization. Cells were stimulated with SARS-CoV-2 S1 and S2 peptide pools and then measured by intracellular staining. (A-B) Percentage of memory $CD4^+$ T cells with (A) $T_H1$ markers (IL-2, TNF or IFNγ) or (B) $T_H2$ markers (IL-4 or IL-13) following stimulation. (C) Percentage of $CD8^+$ T cells expressing IL-2, TNF or IFNγ. (D-E) Percentage of $T_{FH}$ cells that express (D) CD40L or (E) IL-21.

(F-H) BAL fluid was collected at weeks 8, 39 and 43 post-immunization. Lymphocytes in the BAL were stimulated with S1 and S2 peptide pools and responses measured by intracellular cytokine staining using $T_H1$ (F), $T_H2$ (G), and CD8 markers (H). Break in Y-axis indicates a change in scale without a break in the range depicted.

Circles in (A-H) indicate individual NHP. Boxes represent interquartile range with the median denoted by a horizontal line. Dotted lines set at 0%. Reported percentages may be negative due to background subtraction. 8 vaccinated NHP, split into 2 cohorts post-boost.

A

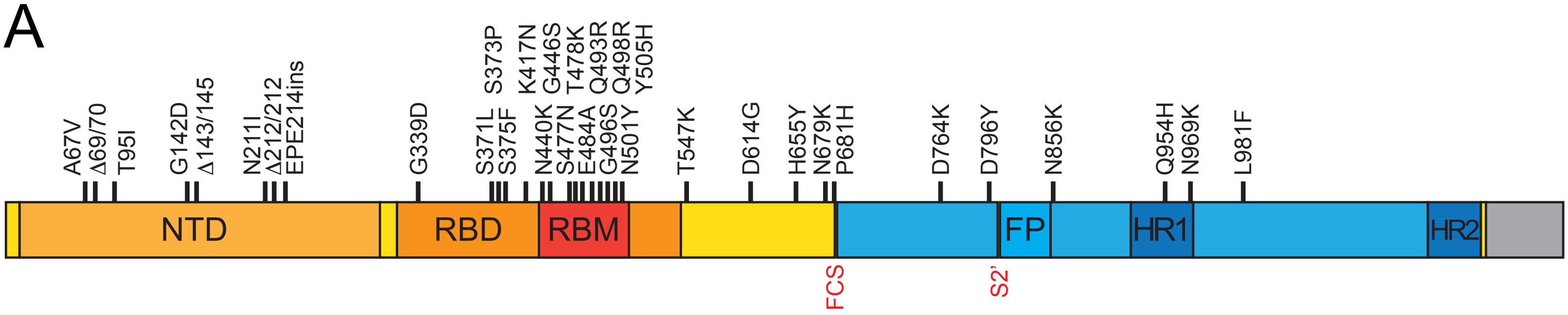


B

| Gene | Amino acid change |
|---|---|
| ORF1ab | K856R |
| ORF1ab | S2083_L2084del insI |
| ORF1ab | A2710T |
| ORF1ab | T3255I |
| ORF1ab | P3395H |
| ORF1ab | L3674_G3676del |
| ORF1ab | I3758V |
| ORF1ab | P4715L |
| ORF1ab | I5967V |
| E | T9I |
| M | D3G |
| M | Q19E |
| M | A63T |
| N | P13L |
| N | E31_S33del |
| N | R203_G204del insKR |

**Figure S7 Omicron challenge stock sequence, related to Figure 5**

(A-B) Omicron stock was sequenced and aligned with Wuhan-Hu-1.

(A) S gene only. Mutations listed above graphic. NTD = N-terminal domain. RBD = receptor binding domain. RBM = receptor binding motif. FP = fusion peptide. HR1 = heptad repeat 1. HR2 = heptad repeat 2. FCS = furin cleavage site. S2' = S2' site.

(B) Whole genome.

| Assay | Primary Regimen | Variant | W6 Reciprocal $ID_{50}$ | W6 Fold change relative to D614G | W41 Reciprocal $ID_{50}$ | W41 Fold change relative to D614G |
|---|---|---|---|---|---|---|
| Live Virus Neutralization | mRNA-1273 | D614G | 6068 | (1.0) | 541 | (1.0) |
| | | Delta | 3790 | 1.6 | 291 | 1.6 |
| | | Beta | 1087 | 7.1 | 148 | 4.8 |
| | | Omicron | 158 | 67.0 | 45 | 12.8 |
| Pseudovirus Neutralization | mRNA-1273 | D614G | 5522 | (1.0) | 324 | (1.0) |
| | | Delta | 2300 | 2.7 | 211 | 1.6 |
| | | Beta | 621 | 13.7 | 148 | 2.9 |
| | | Omicron | 434 | 19.4 | 156 | 2.9 |

| Assay | Second Boost | Variant | W43 Reciprocal $ID_{50}$ | W43 Fold change relative to D614G |
|---|---|---|---|---|
| Live Virus Neutralization | mRNA-1273 | D614G | 9088 | (1.0) |
| | | Delta | 6297 | 1.4 |
| | | Beta | 3122 | 2.9 |
| | | Omicron | 1051 | 8.6 |
| | mRNA-Omicron | D614G | 6069 | (1.0) |
| | | Delta | 4380 | 1.2 |
| | | Beta | 1197 | 5.8 |
| | | Omicron | 718 | 7.9 |
| Pseudovirus Neutralization | mRNA-1273 | D614G | 6670 | (1.0) |
| | | Delta | 3973 | 1.7 |
| | | Beta | 3595 | 2.0 |
| | | Omicron | 4282 | 2.1 |
| | mRNA-Omicron | D614G | 3322 | (1.0) |
| | | Delta | 2244 | 1.3 |
| | | Beta | 1458 | 3.2 |
| | | Omicron | 2039 | 1.5 |

High Low

**Table S1 Reciprocal $ID_{50}$ geometric mean values and fold changes relative to D614G for live virus and pseudovirus neutralization values, related to Figure 1**

NHP were immunized according to Figure S1. The values for reciprocal $ID_{50}$ and fold change relative to D614G at weeks 6 and 41 are averages of 8 animals, while the values at week 43 are the averages of 4 animals. Conditional formatting applied to each individual assay.

| Variant | Mutations outside RBD | Mutations in RBD |
|---|---|---|
| **WA1** | None | None |
| **Beta** | L18F, D80A, D215G, ΔL242-244, R246I, D614G, A701V | K417N, E484K, N501Y |
| **Delta** | T19R, G142D, ΔE156, ΔF157, R158G, D614G, P681R, D950N | L452R, T478K |
| **Omicron** | A67V, ΔH69, ΔV70, T95I, G142D, ΔV143, ΔY144, ΔY145, N211I, L212V_RE, V213P, R214E, T547K, D614G, H655Y, N679K, P681H, N764K, D796Y, N856K, Q954H, N969K, L981F | G339D, S371L, S373P, S375F, K417N, N440K, G446S, S477N, T478K, E484A, Q493R, G496S, Q498R, N501Y, Y505H |

**Table S2 List of mutations in variant-specific S-2P-ACE2 inhibition assays and B cell probes, related to Figure 2**

All variant positions in reference to Wuhan-Hu-1 sequence (Genbank: NC_045512). All S-2P constructs contained furin substitution (GSAS at positions 682-685) and 2P stabilization (K986P and V987P).
